# *In silico* integration of thousands of epigenetic datasets into 707 cell type regulatory annotations improves the trans-ethnic portability of polygenic risk scores

**DOI:** 10.1101/2020.02.21.959510

**Authors:** Tiffany Amariuta, Kazuyoshi Ishigaki, Hiroki Sugishita, Tazro Ohta, Koichi Matsuda, Yoshinori Murakami, Alkes L. Price, Eiryo Kawakami, Chikashi Terao, Soumya Raychaudhuri

**Affiliations:** Center for Data Sciences, Harvard Medical School, Boston, Massachusetts, 02115, USA; Divisions of Genetics and Rheumatology, Department of Medicine, Brigham and Women’s Hospital, Harvard Medical School, Boston, Massachusetts, 02115, USA; Program in Medical and Population Genetics, Broad Institute of MIT and Harvard, Cambridge, Massachusetts, 02142, USA; Department of Biomedical Informatics, Harvard Medical School, Boston, Massachusetts, 02115, USA; Graduate School of Arts and Sciences, Harvard University, Cambridge, Massachusetts, 02138, USA; Laboratory for Statistical and Translational Genetics, RIKEN Center for Integrative Medical Sciences, Kanagawa, 230-0045 Japan; Laboratory for Developmental Genetics, RIKEN Center for Integrative Medical Sciences (IMS), Kanagawa, Japan; Medical Sciences Innovation Hub Program, RIKEN, Kanagawa, Japan; Database Center for Life Science, Joint Support-Center for Data Science Research, Research Organization of Information and Systems, Shizuoka, Japan; Laboratory of Genome Technology, Human Genome Center, Institute of Medical Science, The University of Tokyo, Tokyo, 108-8639 Japan; Laboratory of Clinical Genome Sequencing, Department of Computational Biology and Medical Sciences, Graduate School of Frontier Sciences, The University of Tokyo, Tokyo, 108-8639 Japan; Division of Molecular Pathology, Institute of Medical Science, The University of Tokyo, Tokyo, 108-8639 Japan; Department of Epidemiology, Harvard T.H. Chan School of Public Health, Boston, MA 02115, USA; Department of Biostatistics, Harvard T.H. Chan School of Public Health, Boston, MA 02115, USA; Artificial Intelligence Medicine, Graduate School of Medicine, Chiba University, Chiba, Japan; Clinical Research Center, Shizuoka General Hospital, Shizuoka, 420-8527 Japan; The Department of Applied Genetics, The School of Pharmaceutical Sciences, University of Shizuoka, Shizuoka, 422-8526 Japan; Centre for Genetics and Genomics Versus Arthritis, Centre for Musculoskeletal Research, Manchester Academic Health Science Centre, The University of Manchester, Manchester, UK

## Abstract

Poor trans-ethnic portability of polygenic risk score (PRS) models is a critical issue that may be partially due to limited knowledge of causal variants shared among populations. Hence, leveraging noncoding regulatory annotations that capture genetic variation across populations has the potential to enhance the trans-ethnic portability of PRS. To this end, we constructed a unique resource of 707 cell-type-specific IMPACT regulatory annotations by aggregating 5,345 public epigenetic datasets to predict binding patterns of 142 cell-type-regulating transcription factors across 245 cell types. With this resource, we partitioned the common SNP heritability of diverse polygenic traits and diseases from 111 GWAS summary statistics of European (EUR, average N=180K) and East Asian (EAS, average N=157K) origin. For 95 traits, we were able to identify a single IMPACT annotation most strongly enriched for trait heritability. Across traits, these annotations captured an average of 43.3% of heritability (se = 13.8%) with the top 5% of SNPs. Strikingly, we observed highly concordant polygenic trait regulation between populations: the same regulatory annotations captured statistically indistinguishable SNP heritability (fitted slope = 0.98, se = 0.04). Since IMPACT annotations capture both large and consistent proportions of heritability across populations, prioritizing variants in IMPACT regulatory elements may improve the trans-ethnic portability of PRS. Indeed, we observed that EUR PRS models more accurately predicted 21 tested phenotypes of EAS individuals when variants were prioritized by key IMPACT tracks (49.9% mean relative increase in *R*^2^). Notably, the improvement afforded by IMPACT was greater in the trans-ethnic EUR-to-EAS PRS application than in the EAS-to-EAS application (47.3% vs 20.9%, *P* < 1.7e-4). Overall, our study identifies a crucial role for functional annotations such as IMPACT to improve the trans-ethnic portability of genetic data, and this has important implications for future risk prediction models that work across populations.

## Introduction

An important challenge for complex trait genetics is that there is no clear framework to transfer population-specific genetic data, such as GWAS results, to individuals of other ancestries^1–3^. The importance of this challenge is accentuated by the fact that 80% of all genetic studies have been performed using individuals of European ancestry, accounting for a minority of the world’s population^4^. This is exacerbated by the fact that population-specific linkage disequilibrium (LD) between variants confounds inferences about causal cell types and variants (**Figure 1A**)^5–7^. GWAS have the potential to revolutionize the clinical application and utility of genetic data to the individual, exemplified by current polygenic risk score (PRS) models^5,8–16^.

**Figure 1.**
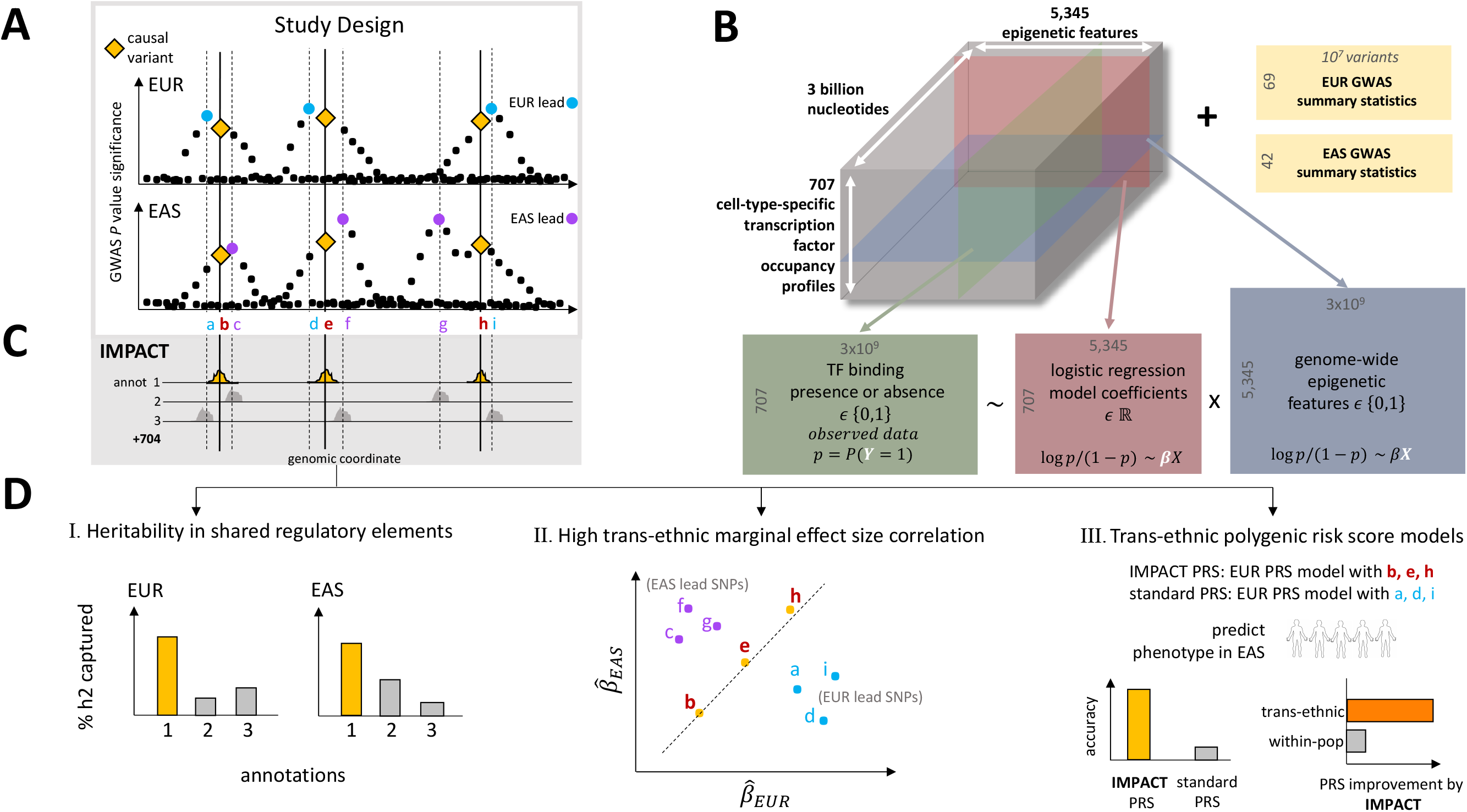
Study design to identify regulatory annotations that prioritize regulatory variants in a multi-ethnic setting. A) Population-specific LD confounding and subsequent inflation of GWAS associations complicate the interpretation of summary statistics and transferability to other populations; functional data may help improve trans-ethnic genetic portability. B) Prism of functional data in IMPACT model: 707 genome-wide TF occupancy profiles (green), 5,345 genome-wide epigenomic feature profiles (blue), and fitted weights for these features (pink) to predict TF binding by logistic regression. Using IMPACT annotations, we investigate 111 GWAS summary datasets (yellow) of EUR and EAS origin. C) Compendium of 707 genome-wide cell-type-specific IMPACT regulatory annotations. D) Annotations that prioritize common regulatory variants must I) capture large proportions of heritability in both populations, II) account for consistent effect size estimations between populations and III) improve the trans-ethnic application of PRS.

However, while the utility of PRS models relies on accurate estimation of allelic effect sizes from GWAS and benefits from genetic similarity between the test cohort and the GWAS cohort, recent studies have explicitly observed a lack of trans-ethnic portability^2,3,5,8,17,18^. Previous studies have extensively shown that functional annotations can improve PRS models when learned and applied to the same population^19,20^, by introducing biologically-relevant priors on causal effect sizes and compensating for inflation of association statistics by LD. However, the potential for functional annotations to improve trans-ethnic PRS frameworks, where the influences of population-specific LD are more profound, has not yet been extensively investigated.

However, designing functional annotations that may improve PRS models is challenging. While the genetic variation of a complex trait likely regulates diverse biological mechanisms genome-wide, such functional annotations must strike a balance of specificity, comprehensively but precisely capturing large regulatory programs. Pinpointing these mechanisms is especially difficult as genome-wide association studies (GWAS) have identified thousands of genetic associations with complex phenotypes^8,21–23^. It has been estimated that about 90% of these associations reside in protein noncoding regions of the genome, making their mechanisms difficult to interpret^24,25^. Defining the etiology of complex traits and diseases requires knowledge of phenotyping-driving cell types in which these associated variants act.

Transcription factors (TFs) are poised to orchestrate large polygenic regulatory programs as genetic variation in their target regions can modulate gene expression, often in cell-type-specific contexts^26,27^. Genomic annotations marking the precise location of TF-mediated cell type regulation can be exploited to elucidate the genetic basis of polygenic traits. However, currently there is no comprehensive catalogue of the binding profiles of the approximately 1,600 human TFs in every known cell type^28^. Moreover, existing TF ChIP-seq datasets are limited to factors with effective antibodies and suffer from inter-experimental variation, noise, and genomic bias^29,30^.

To overcome these challenges, we previously developed IMPACT, a genome-wide cell-type-specific regulatory annotation strategy that models the epigenetic pattern around active TF binding using linear combinations of functional annotations^31^. In rheumatoid arthritis (RA), IMPACT CD4+ T cell annotations captured substantially more heritability than functional annotations derived from single experiments, including TF and histone modification ChIP-seq^6^. In this study, we expanded this approach by aggregating 5,345 functional annotations with IMPACT to create a powerful and generalizable resource of 707 cell-type-specific gene regulatory annotations (**Web Resources**) based on binding profiles of 142 TFs across 245 cell types (**Figure 1B,C**). This study builds on our previous study introducing IMPACT, in which we created only 13 annotations (13 TFs) based on 515 functional annotations. Assuming that causal variants are largely shared between populations^2,21^, we hypothesized that restricting PRS models to variants within trait-relevant IMPACT annotations, which are more likely to have regulatory roles and less likely to be confounded by LD, will especially improve their trans-ethnic portability.

In this study, we identify key IMPACT regulatory annotations that capture genome-wide polygenic mechanisms underlying a diverse set of complex traits, supported by enrichments of genetic heritability, multi-ethnic marginal effect size correlation (a mechanism of improved PRS), and improved trans-ethnic portability of PRS models (**Figure 1D**). Here, we defined and employed our compendium of 707 IMPACT regulatory annotations to study polygenic traits and diseases from 111 GWAS summary datasets of European (EUR) and East Asian (EAS) origin.

Assuming shared causal variants between populations, annotations that prioritize shared regulatory variants must (1) capture disproportionately large amounts of genetic heritability in both populations, (2) be enriched for multi-ethnic marginal effect size correlation, and (3) improve the trans-ethnic applicability of population-specific PRS models. Using our compendium of regulatory annotations, we identified key annotations for each polygenic trait and demonstrated their utility in each of these three applications toward prioritization of shared regulatory variants. Overall, this work improves the interpretation and trans-ethnic portability of genetic data and provides implications for future clinical implementations of risk prediction models.

## Results

### Building a compendium of *in silico* gene regulatory annotations

To capture genetic heritability of diverse polygenic diseases and quantitative traits, we constructed a comprehensive compendium of 707 cell type regulatory annotation tracks. To do this, we applied the IMPACT^31^ framework to 707 unique TF-cell type pairs obtained from a total of 3,181 TF ChIP-seq datasets from NCBI, representing 245 cell types and 142 TFs (**Figure 1B**, **Online Methods**, **Web Resources**, **ST1**, **SF1**)^32^. Briefly, IMPACT learns an epigenetic signature of active TF binding evidenced by ChIP-seq, differentiating bound from unbound TF sequence motifs using logistic regression. We derive this signature from 5,345 epigenetic and sequence features, predominantly generated by ENCODE^33^ and Roadmap^34^ (**Online Methods**, **ST2**); these data were drawn from diverse cell types, representing the biological range of the 707 candidate models. IMPACT then probabilistically annotates the genome, e.g. on a scale from 0 to 1, without using the TF motif, identifying regulatory regions that are similar to those that the TF binds.

To assess the specificity of our IMPACT annotations, we test whether they (1) accurately predict binding of the modeled TF, (2) share cell-type-specific characteristics with other tracks of the same cell type, and (3) score cell-type-specifically expressed genes higher than nonspecific genes. The 707 models that we defined had a high TF binding prediction accuracy with mean AUPRC = 0.74 (se = 0.008, **SF2**) using cross-validation. Annotations segregated by cell type rather than by TF, excluding CTCF, suggesting a single TF may bind to different enhancers in different cell types (**Figure 2A**). Annotations of the same cell types were more strongly correlated genome-wide (Pearson *r* = 0.56, se = 0.06) than annotations of different cell types (Pearson *r* = 0.48, se = 0.003, difference of means *P* < 0.03, **SF2**). Furthermore, the covariance structure between TF ChIP-seq training datasets is similar to that of corresponding IMPACT annotations, indicating that the IMPACT model does not introduce spurious correlations among unrelated ChIP-seq datasets (**SF2**). Lastly, for nine different cell types, we examined cell-type-specifically expressed genes from Finucane et al^35^ and corresponding differential expression *t*-statistics. We observed significantly larger IMPACT probabilities at SNPs in and near these genes (mean = 0.062, se = 0.011) compared to genes that were generally expressed (mean = 0.045, se = 0.006; difference of means *P* = 0.024, **Figure 2B**, **SF2**, **Online Methods**), suggesting that IMPACT annotates relevant cell type regulatory elements.

**Figure 2.**
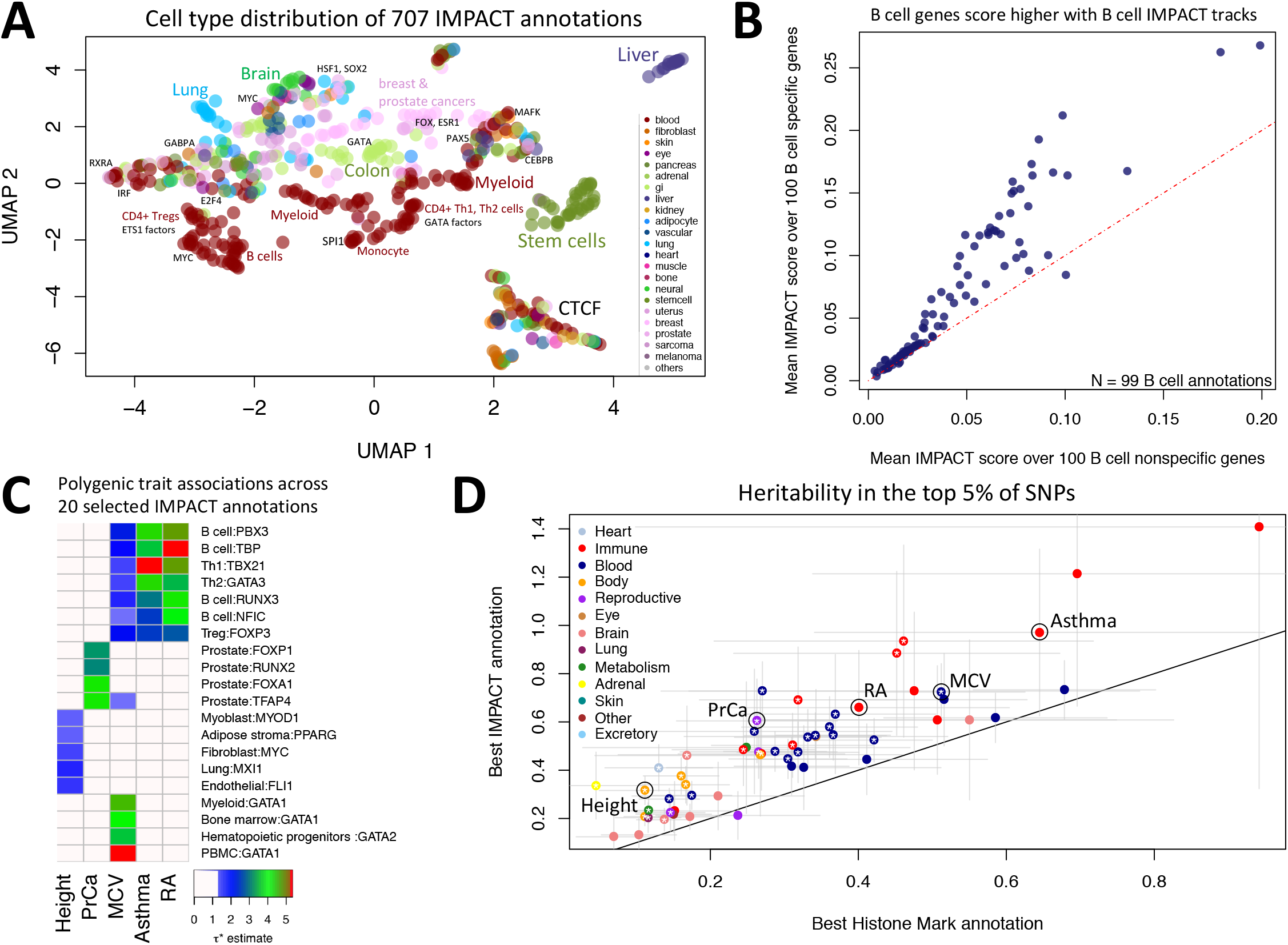
IMPACT annotates relevant cell type regulatory elements. A) Low-dimensional embedding and clustering of 707 IMPACT annotations using uniform manifold approximation projection (UMAP). Annotations colored by cell type category; TF groups indicated where applicable. B) IMPACT annotates cell type specifically expressed genes with higher scores than nonspecific genes. C) Biologically distinct regulatory modules revealed by cell type-trait associations with significantly nonzero τ* across 20 of 707 IMPACT regulatory annotations and 5 representative EUR complex traits, color indicates −log_10_ FDR 5% adjusted *P* value of τ*. D) Lead IMPACT annotations capture more heritability than lead cell-type-specific histone modifications across 60 of 69 EUR summary statistics for which a lead IMPACT annotation was identified. * indicates heritability estimate difference of means *P* < 0.05.

### Partitioning common SNP heritability of 111 GWAS summary statistics in EUR and EAS

We obtained summary statistics from 111 publicly available GWAS summary statistics for diverse polygenic traits and diseases. Throughout the text, we use five randomly selected traits to exemplify our results: asthma, RA, prostate cancer (PrCa), mean corpuscular volume (MCV), and body height. These included 69 from EUR participants^31,36^ (average N = 180K, average heritability z-score = 12.9, 41/69 from UK BioBank)^6,37^ and 42 from EAS participants of BioBank Japan^3,38–40^ (average N = 157K, average heritability z-score = 6.6)^22^ (**ST3**). All of the summary statistics used were generated from studies that had a sample size greater than 10,000 individuals and also had a significantly non-zero heritability (z-score > 1.97). There are 29 phenotypes for which we obtained summary statistics in both EUR and EAS. Although 10/29 traits have a multi-ethnic genetic correlation (*R_g_*) significantly less than 1 (*P* < 0.05/29 tested traits), overall we observed high *R_g_* for most traits, supporting our assumption that causal variants are generally shared across populations (**Online Methods**, **SF3**)^41^. At two extremes, basophil count has a low multi-ethnic *R_g_* of 0.32 (sd = 0.10), while atrial fibrillation has a high multi-ethnic *R_g_* of 0.98 (sd = 0.11), consistent with previous observations made using *Popcorn*, but using different parameter estimation strategies (**Online Methods**)^3^.

We then partitioned the common SNP (minor allele frequency (MAF) > 5%) heritability of these 111 datasets using S-LDSC^6^ with an adapted baseline-LD model excluding cell-type-specific annotations^31,36^ (**SF3, Online Methods**). Next, we tested each of the traits against each of the 707 IMPACT annotations, assessing the significance of a non-zero τ*, which is defined as the proportionate change in per-SNP heritability associated with a one standard deviation increase in the value of the annotation (**Online Methods**)^36^. We observed that 95 phenotypes had at least one significant annotation-trait association (τ* > 0, two-tailed FDR < 5%, **Ext. Data 1, Online Methods**, **ST4-8**). Here, we highlight associations with EUR summary statistics for the five exemplary phenotypes mentioned above: asthma, RA, PrCa, MCV, and height (**Figure 2C**). Consistent with known biology, B and T cells were strongly associated with asthma^42^, RA^43^, and MCV^44,45^ while other blood cell regulatory annotations predominantly derived from GATA factors were also associated with MCV. Prostate cancer cell lines were associated with PrCa, while many cell types including myoblasts^46^, fibroblasts^47^, and adipocytes^48,49^, lung cells, and endothelial cells were associated with height, perhaps related to musculo-skeletal developmental pathways.

For each trait, we defined the lead IMPACT regulatory annotation as the annotation capturing the greatest per-SNP heritability, e.g. the largest while significant τ* estimate (**ST9**). With their top 5% of SNPs, lead IMPACT annotations captured an average of 43.3% heritability (se = 13.8%) across these 95 polygenic traits (**SF4**, **Online Methods**), with more than 25% of heritability captured for two-thirds of the tested summary statistics (73/111 traits) and more than 50% captured for 28% (31/111). Returning to our five exemplary phenotypes, with the top 5% of EUR SNPs, IMPACT captured 97.1% (sd = 17.6%) of asthma heritability with the T-bet Th1 annotation, 65.9% (sd = 12.1%) of RA heritability with the B cell TBP annotation, 60.4% (sd = 8.9%) of PrCa heritability with the prostate cancer cell line (LNCAP) TFAP4 annotation, 72.4% (sd = 6.0%) of MCV heritability with the GATA1 PBMC annotation, and lastly 31.6% (sd = 3.0%) of height heritability with the lung MXI1 annotation (**Figure 2D**). While the observed association between lung and height is not intuitive, within the MXI1 gene lies a genome-wide significant variant associated with height^50^. Moreover, we captured significantly more heritability across EUR traits using our expanded set of 707 IMPACT annotations (mean = 49.5%, se = 12.0%) compared to the 13 annotations in our previous study (mean = 32.3%, se = 1.3%, difference of means *P* = 0.02).

To demonstrate the value of IMPACT tracks, we compared them to annotations derived from single experimental assays. For example, since each of the IMPACT tracks was trained on TF ChIP-seq data, we directly compared the heritability captured by both data types. We observed that the heritability captured by lead IMPACT annotations (mean τ* = 3.53, se = 0.91) was significantly greater than by the analogous TF ChIP-seq used in training (mean τ* = 1.71, se = 0.94, difference of means *P* = 0.02). We also compared IMPACT tracks to histone marks, which are commonly used to quantify cell type heritability^6^. From 220 publicly available cell-type-specific histone mark ChIP-seq annotations of EUR SNPs^6^, we selected the lead histone mark track for each of 69 EUR summary statistics. Restricting to the top 5% of SNPs, we observed that the mean EUR heritability captured by lead IMPACT annotations (49.5%, se = 12.0%) was significantly greater than by lead histone mark annotations (28.4%, se = 9.0%, difference of means *P* = 0.02) (**Figure 2D, ST10**). For example, the lead IMPACT annotation for asthma captured 1.5x more heritability than the best histone mark annotation (H3K27ac in CD4+ Th2), capturing 64.2% (sd = 15.5%) of heritability. Similarly, IMPACT captured 1.7x more RA heritability than H3K4me3 in CD4+ Th17s; IMPACT captured 1.4x more MCV heritability than H3K4me3 in CD34+ cells; IMPACT captured 2.3x more PrCa heritability than H3K4me3 in CD34+ cells; and IMPACT captured 3.1x more height heritability than H3K4me3 in lung cells. In terms of τ*, IMPACT also captured more per-SNP heritability than histone marks: mean τ* fold change = 1.38x (**SF5**).

Since some of our IMPACT annotations are similar to each other (**SF2**), we performed serial conditional analyses in order to identify IMPACT annotations explaining heritability independently from one another (**Online Methods**). This strategy might identify complex traits for which several distinct biological mechanisms are independently regulated by genetic variation. Indeed, we identified 30 EUR phenotypes and 8 EAS phenotypes with multiple independent IMPACT associations (**SF6**, **ST11-12**). For example, four annotations were independently associated with EUR PrCa: prostate (TFAP4), prostate (RUNX2), mesendoderm (PDX1), and cervix (NFYB). Moreover, for seven EUR traits, three IMPACT annotations were independently associated: height (adipocytes, fibroblasts, lung), neutrophil count (monocytes, adipocytes, B cells), osteoporosis (myoblasts, mesenchymal stem cells, cervix), IBD (T cells and two B cell annotations), platelet count (PBMCs, hematopoietic progenitors, muscle), systolic blood pressure (endothelial, mesenchymal stem cells, fibroblasts), and white blood cell count (B cells, adipocytes, hematopoietic progenitors). We found that the heritability z-score, an index correlated with the power of S-LDSC^6^, is strongly predictive of the number of independent regulatory associations (linear regression *P* < 5.4e-9), while sample size is not (linear regression *P* = 0.19) (**SF7**). Our findings suggest that multiple independent regulatory programs can contribute to the heritability of complex traits, and we can detect them when phenotypes are sufficiently heritable and the GWAS provide accurate effect size estimation.

### Concordance of polygenic regulation between European and East Asian populations

Previous studies have shown concordance of polygenic effects between EUR and EAS individuals in RA^1^ and between EUR and African American individuals in PrCa^51^. However, to our knowledge, the extent of these shared effects has not yet been comprehensively investigated across many functional annotations and in diverse traits. Here, we quantified the SNP heritability (τ*) of 29 traits in EUR and EAS captured by a set of approximately 100 independent IMPACT regulatory annotations (**Figure 3B, SF8, Online Methods**). Assuming shared causal variants in EUR and EAS, IMPACT annotations that best prioritize shared genomic regions regulating a phenotype presumably also disproportionately capture similar amounts of heritability in both EUR and EAS (**Figure 1D-I**, **Figure 3A**). Briefly, we selected independent annotations using an iterative pruning approach: for each trait, we ranked all annotations by τ* and removed any annotation correlated with Pearson *r* > 0.5 to the lead annotation and then repeated. As IMPACT annotations are independent of population-specific factors including LD and allele frequencies (**SF3**), they are poised to capture the genome-wide distribution of regulatory variation in a population-independent manner. We observed that τ* estimates across annotations for EUR and EAS are strikingly similar, with a regression coefficient that is consistent with identity (slope = 0.98, se = 0.04). For example, we observed a strong Pearson correlation of τ* between EUR and EAS for asthma (*r* = 0.98), RA (*r* = 0.87), MCV (*r* = 0.96), PrCa (*r* = 0.90), and height (*r* = 0.96). Furthermore, we found that 97.8% of our τ* estimates have no evidence of population heterogeneity (FDR *P* > 0.05) (**Figure 3C**). Among our five representative traits, we observed only one instance of heterogeneity, in which the B cell SRF IMPACT annotation captured RA heritability significantly more in EUR than in EAS (EUR τ* = 1.20 (se = 0.40), EAS τ* = −1.06 (se = 0.46), difference of means *P* < 2.0e-4). Overall, our results suggest that regulatory variants in EUR and EAS populations are equally enriched within the same classes of regulatory elements. This does not exclude the possibility of population-specific variants or causal effect sizes, as evidenced by 10 traits with multi-ethnic genetic correlation significantly less than 1. Rather, these results suggest that causal biology, including disease-driving cell types and their regulatory elements, underlying polygenic traits and diseases, is largely shared between these populations.

**Figure 3.**
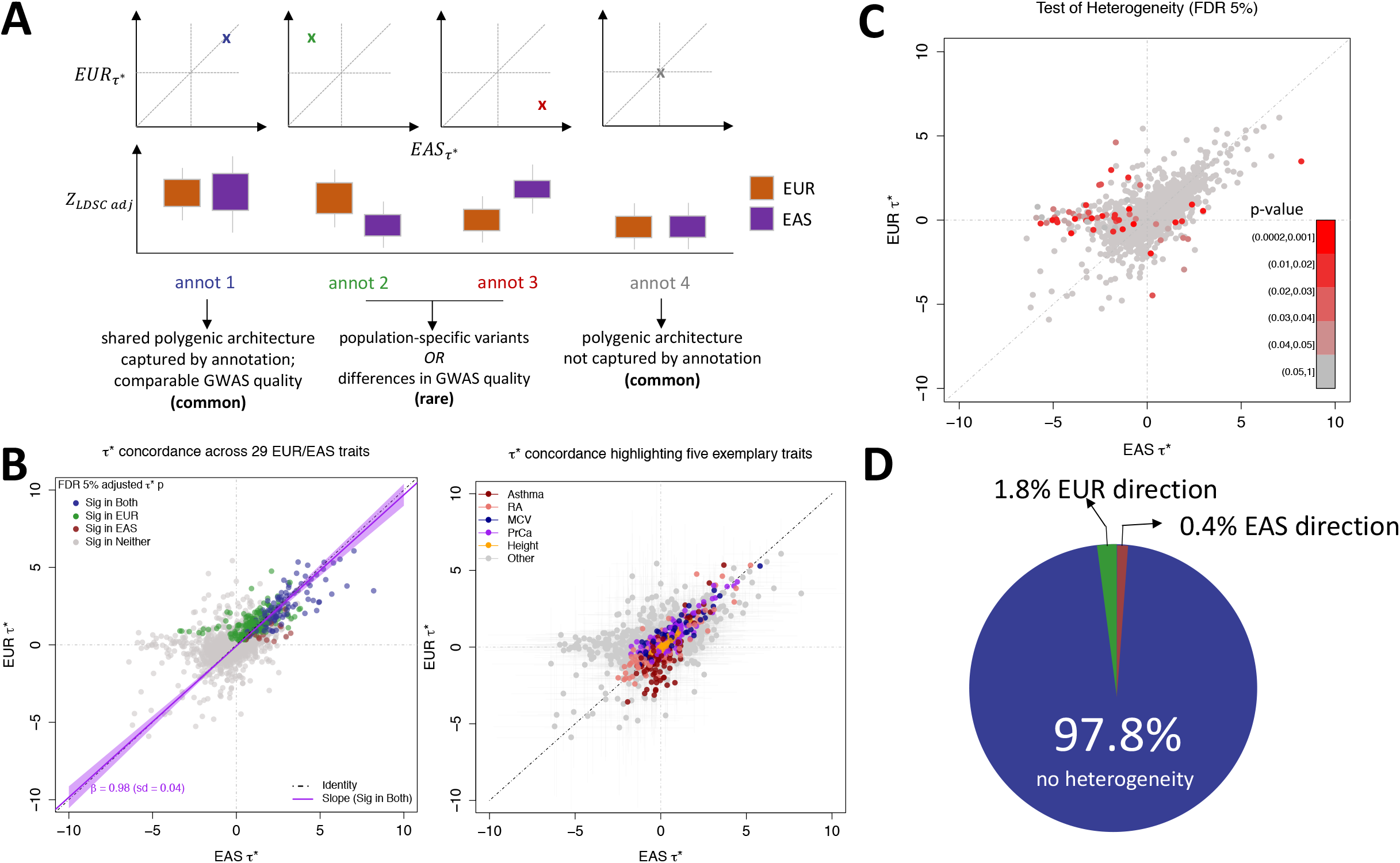
Multi-ethnic concordance of regulatory elements defined by IMPACT. A) Illustrative concept of concordance versus discordance of τ* between populations. Concordance implies a similar distribution of causal variants and effects captured by the same annotation. The implications of discordant τ* are not as straightforward. B) Common per-SNP heritability (τ*) estimate for sets of independent IMPACT annotations across 29 traits shared between EUR and EAS. Left: color indicates τ* significance (τ* greater than 0 at 5% FDR) in both populations (blue), significant in only EUR (green), significant in only EAS (red), significant in neither (gray). Line of best fit through annotations significant in both populations (dark purple line, 95% CI in light purple). Black dotted line is the identity line, y = x. Right: color indicates association to one of five exemplary traits. C) Heterogeneity test at 5% FDR for annotation-trait associations between EUR and EAS. Color indicates significance of difference of means *P* value. D) Heterogeneity test reveals 2.2% of all annotation-trait associations with significantly discordant τ* estimates between populations.

### Models incorporating IMPACT functional annotations improve the trans-ethnic portability of polygenic risk scores

PRS models have great clinical potential: previous studies have shown that individuals with higher PRS have increased risk for disease^8–12^. In the future, polygenic risk assessment may become as common as screening for known mutations of monogenic disease, especially as it has been shown that individuals with severely high PRS may be at similar risk to disease as are carriers of rare monogenic mutations^12^. However, since PRS heavily rely on GWAS with large sample sizes to accurately estimate effect sizes, there is specific demand for the transferability of PRS from populations with larger GWAS to populations underrepresented by GWAS^2,3,5,8,17,18,20^. As we would like to investigate the ability of IMPACT annotations to improve the trans-ethnic application of PRS, we chose pruning and thresholding (P+T) as our model^3,8^. We elected to use P+T rather than LDpred^5,20^ or AnnoPred^19^, which compute a posterior effect size estimate for all SNPs genome-wide based on membership to functional categories. With P+T, we can partition the genome by IMPACT-prioritized and deprioritized SNPs, whereas the assumptions of the LDpred and AnnoPred models do not support the removal of variants, making it difficult to directly assess improvement due to IMPACT prioritization. Moreover, these models have not been explicitly designed or tested for the trans-ethnic application of PRS and thus are beyond the scope of our work. We conventionally define PRS as the product of marginal SNP effect size estimates and imputed allelic dosage (ranging from 0 to 2), summed over M SNPs in the model. Conventional P+T utilizes marginal effect size estimates and therefore is susceptible to selecting a tagging variant over the causal one guided by GWAS *P* values that are inflated by LD. Therefore, we hypothesized that any observed improvement due to incorporation of IMPACT annotations could result from prioritization of variants with higher marginal multi-ethnic effect size correlation (**Figure 1D-II**).

Hence, we tested this hypothesis before assessing PRS performance. We selected 21 of 29 summary statistics shared between EUR and EAS with an identified lead IMPACT association in both populations. Then, using EUR lead IMPACT annotations for each trait (**ST9**), we partitioned the genome three ways: (1) the SNPs within the top 5% of the IMPACT annotation, (2) the SNPs within the bottom 95% of the IMPACT annotation, and (3) the set of all SNPs genome-wide (with no IMPACT prioritization). We then performed stringent LD pruning (&^2^< 0.1 from EUR individuals of phase 3 of 1000 Genomes^52^), guided by the EUR GWAS *P* value, to acquire sets of independent SNPs in order to compute a EUR-EAS marginal effect size estimate correlation (**Online Methods**).

For example, in height, EUR-EAS effect size estimates of SNPs in the top 5% partition (Pearson *r* = 0.434, **Figure 4A**) are 11.4-fold more similar than those in the bottom 95% partition (*r* = 0.038, **Figure 4B**) and 3.31-fold more similar than the set of all SNPs (*r* = 0.131). Meta-analyzed across the 21 traits, the marginal multi-ethnic effect size correlation among the top 5% of IMPACT SNPs was significantly greater than the set of all SNPs genome-wide, across the 10 most lenient of 17 GWAS locus *P* value thresholds examined (all difference of means *P* < 0.026) (**Figure 4C-D**). Furthermore, this observation was consistent across individual traits (**SF9**).

**Figure 4.**
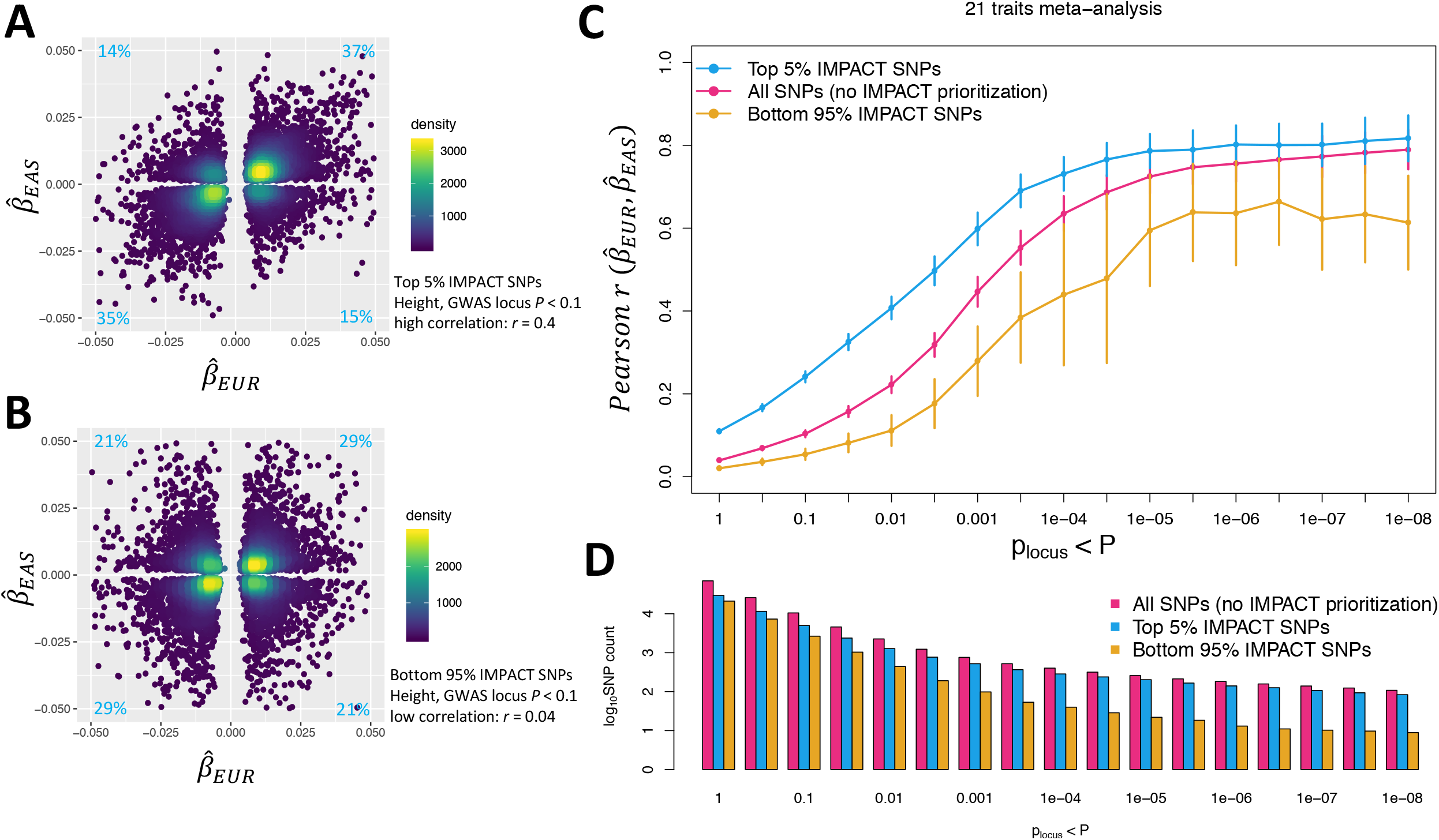
Mechanism by which IMPACT prioritization of shared regulatory variants might improve trans-ethnic PRS performance. A) Estimated effect sizes of variants from genome-wide EUR and EAS height summary statistics in the top 5% of the lead IMPACT annotation for EUR height. Proportions of variants in each quadrant indicated in light blue. B) Estimated effect sizes from genome-wide EUR and EAS height summary statistics of variants in the bottom 95% of the same lead IMPACT annotation for height; mutually exclusive with SNPs in A). C) Meta-analysis of multi-ethnic marginal effect size correlations between populations across 21 traits shared between EUR and EAS cohorts over 17 GWAS *P* value thresholds (with reference to the EUR GWAS). D) Number of SNPs (log10 scale) at each *P* value threshold for each partition of the genome corresponding to C).

For comparison, we performed the same analysis using alternative annotations: lead annotations from 513 cell-type-specifically expressed gene sets (SEG)^35^ and 220 cell-type-specific histone mark annotations (CTS)^6^ (**SF10**). Marginal effect size correlation with IMPACT was comparable to CTS when comparing the top 5% of SNPs to the set of all SNPs (difference of means *P* value > 0.05 at 14 of 17 *P* value thresholds, **SF11**). Compared to SEG, IMPACT-selected SNPs had a significantly greater correlation at 7 of 17 *P* value thresholds (all difference of means *P* value < 0.02, **SF11**). Overall, our results suggest that we might anticipate improved trans-ethnic portability of PRS models by prioritizing SNPs in key IMPACT annotations.

Finally, we addressed our hypothesis that IMPACT annotations improve the trans-ethnic portability of PRS (**Figure 1D-III**). For each of the 21 previously analyzed traits, we built a PRS using effect size estimates from EUR summary statistics and applied it to predict phenotypes of EAS individuals from BioBank Japan (BBJ) (**Figure 5A**). Here, we compare two PRS models, both blind to any EAS genetic or functional information and removing SNPs with LD &*^2^*> 0.2, according to European individuals from phase 3 of 1000 Genomes^52^: (i) standard P+T PRS and (ii) functionally-informed P+T PRS using a subset of SNPs prioritized by the lead EUR IMPACT annotation (**Online Methods**). In functionally-informed PRS models, for each trait separately, we *a priori* selected the subset of top-ranked IMPACT SNPs (top 1%, 5%, 10%, or 50%) which explained the closest to 50% of total trait heritability (**Online Methods**). For all PRS models, we report results from the most accurate model across nine EUR GWAS *P* value thresholds.

**Figure 5.**
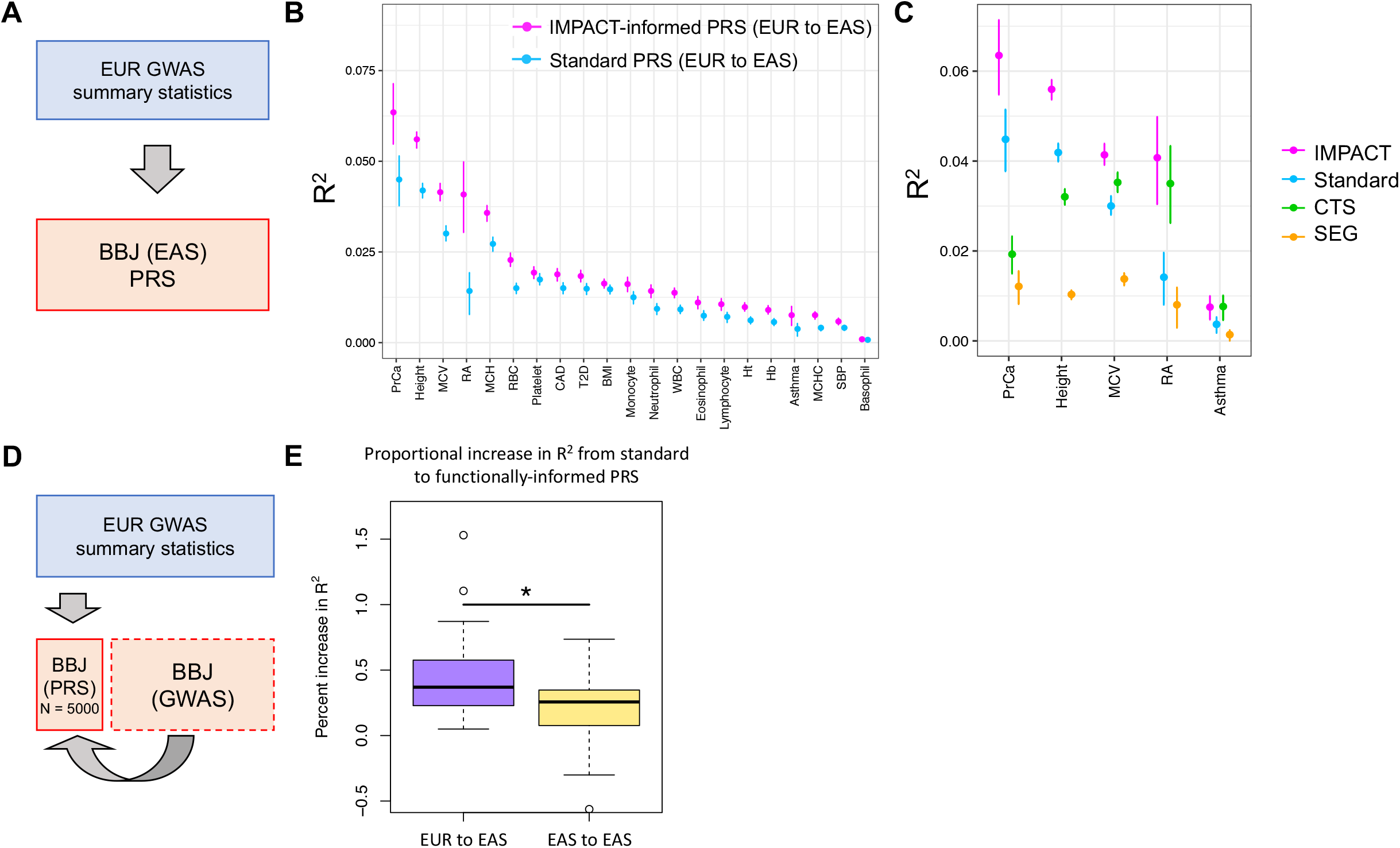
Identifying shared regulatory variants with IMPACT annotations to improve the trans-ethnic portability of PRS. A) Study design applying EUR summary statistics-based PRS models to all individuals in the BBJ cohort. (B) Phenotypic variance (R^2^) of BBJ individuals explained by EUR PRS using two methods: functionally-informed PRS with IMPACT (pink) and standard PRS (blue). Error bars indicate 95% CI calculated via 1,000 bootstraps. C) Phenotypic variance (R^2^) of BBJ individuals across 5 exemplary traits explained by EUR IMPACT annotations relative to lead cell-type-specific histone modification annotations (CTS) and lead cell-type-specifically expressed gene sets (SEG). Error bars indicate 95% CI calculated via 1,000 bootstraps. D) Study design to compare trans-ethnic (EUR to EAS) to within-population (EAS to EAS) improvement afforded by functionally-informed PRS models. For each trait, 5,000 randomly selected individuals from BBJ designated as PRS samples. Remaining BBJ individuals used for GWAS to derive EAS summary statistics-based PRS; no shared individuals between GWAS samples and PRS samples. E) Improvement from standard PRS to functionally-informed PRS compared between trans-ethnic (EUR to EAS) and within-population models (EAS to EAS) using the study design in D).

For each trait, we observed that functionally-informed PRS using IMPACT captured more phenotypic variance than standard PRS (49.9% mean relative increase in *R^2^*, **Figure 5B**, **SF12**, **ST13-15**). The mean phenotypic variance explained across traits by functionally-informed PRS (*R*^2^ = 2.1%, se = 0.2%) was greater than by standard PRS (*R*^2^ = 1.5%, se = 0.1%). For 20 of 21 traits, e.g. excluding basophil count, IMPACT-informed PRS significantly outperformed standard PRS (difference of means *P* < 0.01). Using 10,000 bootstraps of the PRS sample cohort, we found that the IMPACT-informed PRS *R^2^* estimate was consistently greater than the standard PRS estimate for the same 20 traits (all bootstrap *P* < 0.004, **ST15**). We observed the largest improvement for RA from *R^2^*= 1.4% (sd = 0.33%) in the standard PRS versus *R^2^*= 4.1% (sd = %, difference of means *P* < 7.7e-10) in the functionally-informed PRS using the B cell TBP IMPACT annotation. For asthma, *R^2^*= 0.37% (sd = 0.10%) in the standard PRS versus *R^2^*= 0.75% (sd = 0.14%, *P* < 8.5e-4) in the functionally-informed PRS. For MCV, *R^2^*= 3.0% (sd = 0.10%) in the standard PRS versus *R^2^*= 4.1% (sd = 0.12%, *P* < 1.9e-25) in the functionally-informed PRS. For PrCa, *R^2^*= 4.5% (sd = 0.36%) in the standard PRS versus *R^2^*= 6.4% (sd = 0.45%, *P* < 2.4e-6) in the functionally-informed PRS. For height, *R^2^*= 4.2% (sd = 0.10%) in the standard PRS versus *R^2^* = 5.6% (sd = 0.12%, *P* < 1.2e-37) in the functionally-informed PRS.

For our five representative traits asthma, RA, MCV, PrCa, and height, we further compared functionally-informed PRSEUR using IMPACT to models using cell-type-specifically expressed genes (SEG) and cell-type-specific histone modification tracks (CTS)^6,35^ (**Figure 5C**, **ST16**). Across all of the five traits, models using IMPACT explained significantly greater phenotypic variance (mean *R*^2^ = 4.2%, se = 0.3%) than SEG (0.9%, se = 0.1%, all difference of means *P* < 9.9e-11). While IMPACT generally outperformed CTS (*R*^2^ = 2.6%, se = 0.2%, difference of means meta *P* < 1.2e-8), this observation was only individually consistent with 3 of 5 traits (difference of means *P* < 9.3e-8). We performed a similar bootstrap analysis as above, yielding similar results; for only RA and asthma did IMPACT-PRS not produce consistently greater *R^2^* estimates than CTS-PRS (**ST16**).

Functionally-informed PRS might to some extent compensate for population-specific LD differences between populations. Hence, we hypothesized that IMPACT-informed PRS would improve standard PRS moreso in the trans-ethnic prediction framework, in which EUR PRS models predict EAS phenotypes, than in a within-population framework, in which EAS PRS models predict EAS phenotypes. Here, we define within-population PRS as PRSeas and trans-ethnic PRS as PRSeur to avoid confusion. In order to directly compare PRS model improvements between PRSeas and PRSeur, we evaluated prediction accuracy on the same individuals. Briefly, we partitioned the BBJ cohort to reserve 5,000 individuals for PRS testing, derived GWAS summary statistics from the remaining individuals, and performed P+T PRS modeling and prediction as done above (**Figure 5D, SF13-15, ST17-18, Online Methods**). For functionally-informed PRSeas, we selected lead IMPACT annotations from S-LDSC results using GWAS summary statistics, as done above, on the partition of the BBJ cohort excluding the 5,000 PRS test individuals. We defined improvement as the percent increase in *R*^2^ from standard to functionally-informed PRS; therefore, differences in PRS performance due to intrinsic factors, such as GWAS power or genotyping platform, cancel out. In both scenarios, we observed significant non-zero improvements: averaged across the 21 traits in the trans-ethnic setting (mean percent increase in *R*^2^ = 47.3%, se = 8.1%, *P* < 2.7e-9) and in the within-population setting (mean percent increase in *R*^2^ = 20.9%, se = 6.6%, *P* < 7.5e-4). Indeed, this revealed a significantly greater improvement in the trans-ethnic than in the within-population application (difference of means *P* < 1.7e-4, **Figure 5E**).

Overall, our results reveal that functional prioritization of SNPs using IMPACT significantly improves both trans-ethnic and within-population PRS models, but is especially advantageous for the trans-ethnic application of PRS. In conclusion, our results nominate the prioritization of SNPs according to functional annotations, especially using IMPACT, as a potential tentative solution for the lack of trans-ethnic portability of PRS models. While individuals of European ancestry dominate current genetic studies, population-nonspecific cell-type-specific IMPACT annotations can help transfer highly powered EUR genetic data to study still underserved populations.

## Discussion

In this study, we created a compendium of 707 cell-type-specific regulatory annotations (**Web Resources**) capturing disproportionately large amounts of polygenic heritability in 95 complex traits and diseases in EUR and EAS populations. We then proposed a three-step framework to assess how well prioritization of regulatory variants with functional data can improve multi-ethnic genetic comparisons. First, we showed that heritability-enriched regulatory elements between EUR and EAS populations capture indistinguishable proportions of heritability across 29 complex traits. Second, we showed that functional prioritization of variants selects those with more highly correlated marginal effect sizes between populations; this might explain the improvement driven by functional prioritization in P+T PRS models which use marginal effect sizes. Third, we showed that variant prioritization with IMPACT annotations results in consistently improved PRS prediction accuracy, especially for the trans-ethnic application; potentially due to overcoming large population-specific influences such as LD, an important challenge of multi-population models.

Designing genetic models for each complex trait or disease that capture risk for the full diversity of the human population will be challenging. This necessitates approaches that effectively transfer predictive genetic information from well studied populations to less well studied populations. Our work provides insight into the potential clinical implementation of PRS and broader genetic applications that aim to integrate multi-ethnic data. We argue for the use of biologically diverse IMPACT annotations to capture relevant genetic signal and compensate, to some extent, for differences in LD across populations. While we did not assess a PRS model using meta-analyzed summary statistics from two or more populations in this study, we believe that this approach could be effective, especially for populations with limited GWAS sample size.

We believe that IMPACT may prioritize phenotype-driving regulatory variation. We have shown IMPACT to be more effective at capturing genetic variation of complex traits than commonly used functional annotations such as experimentally-derived cell-type-specific histone marks or gene sets. We hypothesize the utility of IMPACT comes from 1) cell-type-specificity of TF binding models which locate key classes of regulatory elements and 2) the integration of thousands of experimentally-derived annotations, which presumably removes noise and enriches for biological signal present in each individual annotation. Here, we did not demonstrate the potential utility of IMPACT to perform functional fine-mapping to reduce credible sets beyond our previous work^31^, due to lack of sufficient gold standards with causal experimental validation and the limitation to genome-wide significant variants. The specific application of IMPACT in multi-ethnic fine-mapping needs to be further investigated.

We must consider several important limitations of our work. First, our functional insights are limited to cell types with publicly available TF ChIP-seq data, lacking ones that are rarer or more difficult to assay. In the future, the cell-type-specific functional training data for IMPACT may be replaced by newer experimental strategies to map enhancers. For example, high-throughput CRISPR screens paired with assays for open chromatin could be used to precisely redefine the regulatory landscape. Second, we used multi-ethnic data to argue for the utility of our approach. However, the robustness of multi-ethnic comparisons for a given phenotype rely on properties surrounding the recruitment of individuals or the exact genotyping platform used in various biobanks, which may result in cohort-bias that inflates within-population PRS prediction accuracy. For example, BBJ is a disease ascertainment cohort, in which each individual has any one of 47 common diseases^53,54^; therefore, BBJ control samples are not comparable to healthy controls of UKBB. Other biases may arise from clinical differences in phenotyping. Also, we only considered a single non-EUR population in this study, while the disparity in trans-ethnic portability and hence resulting benefit from functional annotations may be greater in other non-EUR populations.

In conclusion, we demonstrated that IMPACT annotations improve the comparison of genetic data between populations and trans-ethnic portability of PRS models using ancestrally unmatched data. While a long-term goal of the field must be to diversify GWAS and other genetic studies in non-European populations, it is imperative that genetic models be developed that work in multiple populations. Such initiatives will necessitate the use of population-independent functional annotations, such as IMPACT, in order to capture shared biological mechanisms regulated by complex genetic variation.

## Supplemental Data

See *Supplement.pdf* and *Supplementary_Tables.xlsx*

## Online Methods

See *Online_Methods.pdf*

## Acknowledgements

This work is supported in part by funding from the National Institutes of Health (NHGRI T32 HG002295, UH2AR067677, 1U01HG009088, U01 HG009379, and 1R01AR063759) and the Doris Duke Charitable Foundation grant #2013097.

## Web Resources

1. IMPACT 707 annotations: https://github.com/immunogenomics/IMPACT/tree/master/IMPACT707
2. IMPACT Github repository: https://github.com/immunogenomics/IMPACT
3. HOMER: http://homer.ucsd.edu/homer/motif/
4. S-LDSC: https://github.com/bulik/ldsc
5. 1000 Genomes: http://www.internationalgenome.org/
6. Cell-type-specifically expressed gene set annotations and LD scores: https://data.broadinstitute.org/alkesgroup/LDSCORE/LDSC_SEG_ldscores/
7. Cell-type-specific histone modification ChIP-seq datasets: https://data.broadinstitute.org/alkesgroup/LDSCORE/
8. Plink: https://www.cog-genomics.org/plink2

## Online Methods

### Data

#### TF ChIP-seq data

We previously collected 3,181 publicly available transcription factor (TF) chromatin immunoprecipitation (ChIP) datasets derived from human primary cells or cell lines. We downloaded raw sequencing data in SRA format from NCBI GEO, then converted the data to FASTQ format using the SRA Toolkit function fastq-dump, used FastQC for quality assessment of sequencing reads, and finally mapped reads to the human genome (hg19/GRCh37) with Bowtie2 [v2.2.5] using default parameters. All ChIP-seq datasets were matched to corresponding control data from which peaks were called with macs [v2.1] with q value < 0.01 under a bimodal model, producing 3,181 bed file-formatted files^1,2^. The 1,542 datasets selected for use with our IMPACT model framework (see below) are listed with accession codes in **ST1**.

#### Genome-wide annotation data

We augmented our set of 515 publicly available epigenomic and sequence feature annotations from our previous study^3^ with 116 personally curated datasets from NCBI, 2,593 ENCODE histone ChIP-seq datasets and 2,121 ENCODE open chromatin DNase-seq datasets^4^, all publicly available at the accessions provided in **ST2**. All files were collected in 6-column standard bed file format. This augmentation brought the total number of features to 5,345.

#### Genome-wide association data

We collected publicly available summary statistics data for 111 genome-wide association studies (GWAS) across separate cohorts of East Asian and European individuals^5–7^. East Asian GWAS data were collected from Biobank Japan (BBJ) while European GWAS data were collected from either UKBioBank (UKBB) or the GWAS catalog, referred to as PASS (publicly available summary statistics) (**ST3**). All GWAS summary statistics were reformatted to be compatible with S-LDSC (see below) and thus contained the following information for each SNP (per row): rsID, A1 (reference allele), A2 (alternative allele), GWAS sample size (effective sample size per SNP, may vary with genotyping), chi-square statistic, *z*-score. For multi-ethnic genetic correlation and polygenic risk score prediction, all GWAS summary statistics were reformatted to contain the SNP ID (chr_position_A1_A2), chromosome, base pair, A1, A2, effect size estimate, effect size estimate standard error, and *P*-value.

#### Cell-type-specifically expressed gene set (SEG) and cell-type-specific histone modification (CTS) annotations

We downloaded 513 publicly available SEG annotations for European SNPs from phase 3 of 1000 Genomes accompanied by pre-computed LD scores (see **Web Resources**)^8^. SEG annotations are binary and thus each SNP is designated a 1 or a 0, indicating that the SNP does or does not lie, respectively, within 100 kb of the gene body of the corresponding gene set^8^. We downloaded 220 publicly available CTS annotations of peak data in bed file format, from which we annotated European SNPs from phase 3 of 1000 Genomes^9^ and used S-LDSC to compute LD scores (see **Web Resources**)^7^. These annotations are also binary, in which case each SNP is designated a 1 or a 0, indicating that the SNP does or does not like, respectively, within the peak of histone modification.

#### BioBank Japan data

For PRS analysis, we utilized phenotype and genotype data of the BioBank Japan Project (BBJ)^10,11^. All of the calculations related to PRS were conducted on the RIKEN computing server. BBJ is a biobank that collaboratively collects DNA and serum samples from 12 medical institutions in Japan. This project recruited approximately 200,000 patients with the diagnosis of at least one of 47 diseases. Informed consent was obtained from all participants by following the protocols approved by their institutional ethical committees. We obtained approval from the ethics committees of the RIKEN Center for Integrative Medical Sciences and the Institute of Medical Sciences at the University of Tokyo.

### Statistical Methods

#### IMPACT Model

We implemented our previously defined model to predict TF binding on a motif site. This model regresses the likelihood (*p*) of a binding event on the epigenomic profile of the motif site, in a logistic regression framework over *j* epigenomic features as follows:

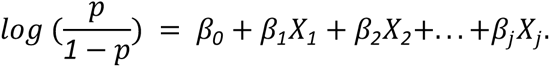

We use a weighted average of ridge and lasso regularization terms in the objective function to restrict the magnitude of fit coefficients and enforce sparsity to reduce overfitting, respectively, as follows:

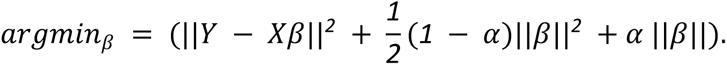

#### Training IMPACT

We trained an IMPACT model for each unique cell type-TF pair present in our data collection. Our collection consists of 3,181 TF ChIP-seq profiles, representing 442 TFs, 296 cell types, and 24 tissues. The IMPACT model requires that the assayed TF has a distinct binding motif and so we removed all ChIP-seq datasets corresponding to a TF that did not have a known sequence motif in MEME, Jaspar, or Transfac databases. This resulted in 1,542 TF ChIP-seq profiles across 142 TFs, 245 cell types, 23 tissues, and 728 unique combinations of TFs and cell types. As we did in our previous study^3^, we merged experiments of the same TF-cell type combination by taking the union of the peaks. We next identified motif sites bound by a TF by using HOMER [v4.8.3]^12^ to scan ChIP-seq peaks for motif matches exceeding the empirically determined motif detection threshold. Similarly, we identified motif sites not bound by a TF by using HOMER to scan the entire genome for sequence matches. 21 of these models did not contain sufficient overlap between TF sequence motifs and ChIP-seq peaks which would lead to underfitting in the logistic regression (fewer than 7), thereby resulting in 707 total possible IMPACT annotations. We then trained 707 IMPACT models using up to 1,000 TF-bound sequence motifs (evidenced by ChIP-seq) and 10,000 unbound sequence motifs. For each of 707 TF-cell type pairs, we learned a predictive model of TF binding and annotated SNPs genome-wide for both EUR and EAS populations, with a mean regulatory probability per nucleotide of 0.02 (se = 7.5e-4).

#### Assessing cell type specificity of IMPACT tracks

We acquired lists of specifically expressed genes in 9 different cell types: T cells, B cells, fibroblasts, monocytes, brain, liver, colon, prostate, and breast according to differential gene expression *t*-statistics from previous work^8^, specifically labeled as T.4+8int.Th, B.Fo.LN, Cells_Transformed_fibroblasts, Mo.6C+II-.LN, Brain_Cortex, Liver, Colon_Transverse, Prostate, Breast_Mammary_Tissue, respectively from either ImmGen or GTEx databases. Large and positive *t*-statistics represent greater specificity of gene expression in the target cell type, large but negative *t*-statistics represent specifically repressed genes, and *t*-statistics near 0 represent nonspecific gene expression, representing commonly expressed genes. For each cell type, we selected the 100 genes with highest *t*-statistics, e.g. specifically expressed (SE) genes, and 100 genes such that −0.5 < *t*-statistic < 0.5, e.g. not specifically expressed genes (NS). For each cell type separately, we collected all related IMPACT annotations from the compendium of 707 total annotations. Then for each annotation separately, we computed the average IMPACT score over all EUR SNPs from phase 3 of 1000 Genomes within 2kb of each SE or NS gene body. Finally, we computed the average across all 100 SE and 100 NS genes, separately.

#### Partitioning heritability with S-LDSC

We applied S-LDSC [v1.0.0]^7^ to partition the common (MAF > 5%) SNP heritability of 111 polygenic traits and diseases, with significantly non-zero heritability estimates (*P* < 0.05). We partitioned heritability using a customized version of the baselineLD model, in which we excluded cell-type-specific regulatory annotations (as we would be testing the enrichment of such annotations from IMPACT). In total, we used 69 cell-type-nonspecific baselineLD annotations and added one or more IMPACT annotations to the model to test for cell-type-specific enrichment. We use three metrics to evaluate how well our IMPACT annotations capture polygenic heritability: enrichment^7^, the proportion of heritability explained by the top 5% of SNPs^7^, and per-annotation standardized effect size, τ*^6^. Briefly, enrichment is defined as the proportion of common SNP heritability divided by the genome-wide proportion of SNPs in the annotation, for continuous annotations this is the average annotation value across SNPs. τ* represents the average per-SNP heritability of a category of SNPs, where a single SNP may claim membership to one or more categories. τ* is defined as the proportionate change in per-SNP heritability associated with a one standard deviation increase in the value of the annotation. The sum of the τ* over categories of SNPs equals the total estimated heritability of the trait. τ* has units of heritability and is comparable between traits, annotations, and populations, because it is normalized for the total heritability (indicative of the power of the GWAS), the dispersion of the annotation values (annotation size), and the number of common SNPs (population-specific) considered in the model, respectively. 7, the precursor of τ*, is the coefficient estimated in the S-LDSC regression. 7 and τ* are conditionally dependent on the provided baselineLD annotations. Therefore, the τ* estimate for an IMPACT annotation is considered a measure of cell-type-specific or annotation-specific SNP heritability, as the remaining annotations in the model (baselineLD) are not cell-type-specific. Significance of τ* is computed using a *z*-test of how different the τ* estimate is from 0; the significance of strictly positive τ* estimates are reported in our study.

#### Measuring heritability in top X% of SNPs of a continuous annotation

To partition the heritability captured by various top echelons of SNPs of a given continuous annotation, we used the same strategy as in a previous study^6^. By this strategy, the proportion of heritability explained by a set of SNPs is the sum over all SNPs of the product of the τ* of each category in the S-LDSC model, e.g. baselineLD plus IMPACT annotation, and the SNP membership to that category (1 or 0 in the case of binary annotations, continuous values in the case of continuous annotations) divided by the same metric for all SNPs genome-wide.

#### Conditional S-LDSC analysis to identify independent annotation-trait associations

Due to the redundancy in modeled cell type programs and inherent covariance of IMPACT annotations (**SF2**), the τ* associations we find with S-LDSC cannot be independent. To this end, for each of 95 traits across EUR and EAS for which we identified a lead IMPACT annotation, reported in **ST9**, we performed a series of conditional analyses using S-LDSC. For each trait with more than one significant τ* association, we created S-LDSC models consisting of the 69 baselineLD annotations, the lead annotation for that trait, and separately, each remaining significant IMPACT annotation. We kept annotations that retained their τ* significance when conditioned on the lead annotation(s), which we also required to retain significance. We iteratively performed these conditional analyses until we were no longer able to identify independent τ* associations.

#### Deming regression of EUR τ* on EAS τ*

As there is significant correlation among IMPACT annotations, due to redundancy in cell type regulatory elements, we used an iterative pruning approach, similar to LD-pruning, to identify independent IMPACT annotations. For each trait, we ranked all 707 IMPACT annotations by their τ* significance values. Then, we selected the lead annotation, removed all annotations correlated with Pearson *r* > 0.5, and selected the next lead annotation, and so on. This approach produced a set of relatively independent annotations, for which the assumptions of Deming, or any, regression would not be violated.

For each trait, we ran Deming regression over approximately 100 independent IMPACT annotations using the R function *deming* within the package *deming*. Across independent observations for all traits, we tested the null hypothesis that the slope of the Deming regression, which considers standard errors on both the predictor (EUR τ*) and response variables (EAS τ*), is equal to 1.

#### Multi-ethnic and within-population genetic correlation

We computed the genetic correlation (*R_g_*) between pairs of 29 traits for which we acquired EUR and EAS GWAS using Popcorn [v.0.9.6]^13^ with default parameters, including maximum likelihood estimation as opposed to regression^14^. First, we computed cross-population scores between the two populations using the *compute* flag with the *popcorn* executable, indicating approximately the correlation between LD at each SNP using EUR and EAS reference LD panels from phase 3 of 1000 Genomes. Then, we used the *fit* flag with the *popcorn* executable to compute the multi-ethnic genetic correlation of these 29 traits. *R_g_* estimates computed after restricting to MAF > 5% did not significantly differ from no MAF restriction. Popcorn computes *R_g_* using either “genetic impact” (effect sizes normalized by allele frequency) or “genetic effect” (unmodified effect sizes). We observed no significant heterogeneity between the *R_g_* computed using “genetic impact” and “effect”, although “genetic effect” estimates were consistently but not significantly larger.

We then computed cross-trait cross-population genetic correlations across 21 traits for which we observed at least one significant IMPACT annotation association in both EUR and EAS. Therefore, in total we computed the genetic correlation among 42 traits (21 phenotypes x 2 populations). For pairs of traits with one from EUR and one from EAS, we used Popcorn as described above with MAF threshold of 5% and “genetic impact”. For pairs of traits from the same population we used LDSC [v.1.0.0]. First we used the *munge_sumstats.py* script to make the direction of allelic effect consistent in the GWAS summary statistics while also restricting to well-imputed Hapmap3 SNPs. Then, we used the *ldsc.py* script with the *-rg* flag to compute the genetic correlation using EUR and EAS reference LD panels from phase 3 of 1000 Genomes where appropriate.

#### Multi-ethnic marginal effect size correlation

We acquired GWAS summary statistics for each of 21 shared traits between EUR and EAS for which there was at least one significant IMPACT association in each population. Then, we restricted to SNPs shared between EUR and EAS GWAS summary statistics. Next, we performed stringent iterative LD clumping with PLINK [v1.90b3]^15^ using EUR summary statistics (selecting the most significant SNP, then removing all SNPs in LD with *r^2^* > 0.1 within 1 Mb, then selecting the next most significant SNP, and so on).

This step satisfies the assumption of independence in the Pearson correlation that we will compute among marginal effect sizes. We selected our initial set of SNPs under three scenarios: (1) using no functional inference, (2) using the top 5% of SNPs according to the trait’s lead EUR IMPACT annotation, and (3) using the bottom 95% of SNPs according to the trait’s lead EUR IMPACT annotation (mutually exclusive with scenario 2). With our set of independent SNPs for each trait and under each of three scenarios, we compute a Pearson correlation between the estimated effect sizes, while further stratifying loci on 17 EUR *P*-values (1, 0.3, 0.1, 0.03, 0.01, 3e-3, 1e-3, 3e-4, 1e-4, 3e-5, 1e-5, 3e-6, 1e-6, 3e-7, 1e-7, 3e-8, 1e-8). For example, stratum with *P* = 0.1 includes all SNPs with EUR GWAS *P* < 0.1.

#### Polygenic risk score calculation

In this study, we utilized pruning and thresholding (P+T) for the calculation of PRS. We constructed PRS models from either EUR summary statistics or EAS summary statistics and evaluated their predictive performance on individual EAS phenotypes.

Here, we define within-population PRS as PRSeas and trans-ethnic PRS as PRSeur to avoid confusion. For PRSeur, we utilized genome-wide summary statistics from EUR as reported in their publicly available version. For PRSeas, we held out 5,000 individuals for PRS analysis and conducted GWAS using the remaining individuals to avoid overfitting (see next section). For each trait separately, we restricted our analysis to variants that exist in both GWAS summary statistics and post-imputation genotype data of EAS individuals used for PRS analysis (imputation quality of *r^2^* > 0.3 in minimac3). A detailed description related to the genotyping platform and imputation strategy is provided in a previous report^2^. We excluded the MHC region in this analysis.

We designed PRS models using two strategies: standard PRS and functionally-informed PRS. For standard PRSeur, we performed conventional LD clumping to acquire sets of independent SNPs using EUR LD reference panels from phase3 of 1000 Genomes. Similarly for PRSeas, we utilized EAS LD reference panels from phase3 of 1000 Genomes. We used PLINK [v1.90b3]^15^ to remove variants in LD with *r^2^* > 0.2 with a significance threshold for index SNPs of *P* = 0.5. For functionally-informed PRS, we restricted the analysis to variants with high IMPACT score according to the lead IMPACT annotation before conducting LD clumping. As before, we define the lead annotation as the one with the largest τ* estimate that was significantly greater than 0. When we designed PRSeur, we utilized the lead IMPACT annotation in EUR GWAS summary statistics (EAS summary statistics were not taken into account to avoid overfitting). Similarly, when we design PRSeas, we utilized the lead IMPACT annotation in EAS GWAS summary statistics for which 5,000 EAS individuals for PRS analysis were removed to avoid overfitting. We performed LD clumping using variants within a predefined top percentage of IMPACT scores. This was determined by the percentage that captured the closest to 50% of total trait heritability; considered percentages included the top 1%, 5%, 10%, and 50%.

We evaluated PRS performance using EAS individuals. First, we used all individuals in the BBJ cohort for PRSeur testing. Second, we compared the improvement afforded by IMPACT in PRSeur relative to PRSeas models using 5,000 randomly selected individuals in BBJ; specifically for case-control GWAS, we randomly selected 1,000 cases and 4,000 controls.

For all models, we built a PRS for each individual *j* in our test set (in all cases, there is no overlap between GWAS samples and PRS samples) using variant effect size estimates from GWAS as follows:

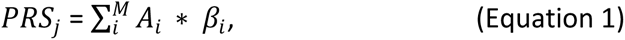

Where M is the total number of SNPs shared between GWAS summary statistics and post-imputation genotype data of EAS individuals, *i* is the *i*^th^ SNP in the model, *A*_*i*_ is the allele dosage of the trait-increasing allele *i*, and *β*_*i*_ is the estimated effect size of allele *i* from the GWAS. We calculated PRS using PLINK2.

For QC of quantitative phenotypes, we excluded (1) related samples (PI_HAT > 0.187 estimated by PLINK), (2) samples with age < 18 and age > 85, and (3) samples with measured values outside three interquartile ranges (IQR) of the upper or lower quartiles. The effect of sex, age, *age*^2^, the top 10 PCs, and affection status of 47 diseases were removed by linear regression, and the residuals were further normalized by the rank-based inverse normal transformation (see Equation 3 below). For QC of case/control phenotypes, we excluded (1) related samples (PI_HAT > 0.187 estimated by PLINK) and (2) samples with age < 18 and age > 85.

We then regressed our phenotype of interest (Y), a measured quantitative trait or a diagnosed disease among the PRS samples, on the per-individual PRS as follows:

For diseases,

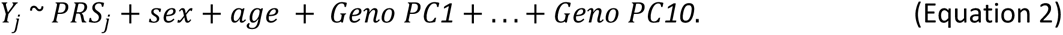

For quantitative traits,

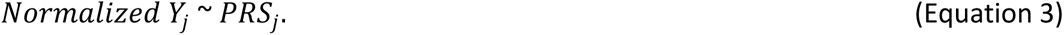

We then report the variance explained; for quantitative traits, this is the variance explained by a linear model and for diseases, the variance explained is from a logistic model (Nagelkerke *R^2^*)^14,16,17^ which we convert to liability scale pseudo *R^2^* such that *R^2^* values are comparable among both quantitative and case/control phenotypes. We used various GWAS *P* value thresholds (0.1, 0.03, 0.01, 0.003, 0.001, 3e-4, 1e-4, 3e-5, 1e-5) to assess the predictive performance of our PRS. For each model, we reported in the text the largest *R^2^*achieved across the nine P value thresholds.

To estimate confidence intervals of PRS performance (*R^2^*, as explained above), we conducted 1,000 bootstraps using the R package *boot*. We also conducted 10,000 bootstraps to evaluate whether the *R^2^*difference between two PRS models (functionally-informed - standard) is significantly greater than 0; we calculated the *R^2^*difference between two PRS models in each round of bootstrapping (delta *R^2^*), and assess its distribution in 10,000 bootstraps. If we let N be the frequency of delta *R^2^*< 0, we define one-sided *P* values for delta *R^2^*> 0 as (N + 1)/10,000.

### Genome-wide association studies in BBJ

As described in the previous section, we held out 5,000 randomly selected individuals for the PRS analysis and performed GWAS on the remaining individuals (sample sizes are provided in **ST13-14**). GWAS was conducted with PLINK2 using the same imputed dosages as used in the PRS analysis. For quantitative traits, normalized residuals were analyzed by a linear regression model. For diseases, affection status was analyzed by a logistic regression model using age, sex, and the top 10 genotype PCs as covariates.

## References

1. Kichaev, G. & Pasaniuc, B. Leveraging Functional-Annotation Data in Trans-ethnic Fine-Mapping Studies. Am. J. Hum. Genet. 97, 260–271 (2015).

2. Lam, M. et al. Comparative genetic architectures of schizophrenia in East Asian and European populations. doi:10.1101/445874

3. Martin, A. R. et al. Clinical use of current polygenic risk scores may exacerbate health disparities. Nat. Genet. 51, 584–591 (2019).

4. Sirugo, G., Williams, S. M. & Tishkoff, S. A. The Missing Diversity in Human Genetic Studies. Cell 177, 26–31 (2019).

5. Vilhjálmsson, B. J. et al. Modeling Linkage Disequilibrium Increases Accuracy of Polygenic Risk Scores. Am. J. Hum. Genet. 97, 576–592 (2015).

6. Finucane, H. K. et al. Partitioning heritability by functional annotation using genome-wide association summary statistics. Nat. Genet. 47, 1228–1235 (2015).

7. Bulik-Sullivan, B. K. et al. LD Score regression distinguishes confounding from polygenicity in genome-wide association studies. Nat. Genet. 47, 291–295 (2015).

8. International Schizophrenia Consortium et al. Common polygenic variation contributes to risk of schizophrenia and bipolar disorder. Nature 460, 748–752 (2009).

9. Chatterjee, N., Shi, J. & García-Closas, M. Developing and evaluating polygenic risk prediction models for stratified disease prevention. Nat. Rev. Genet. 17, 392–406 (2016).

10. Stahl, E. A. et al. Bayesian inference analyses of the polygenic architecture of rheumatoid arthritis. Nat. Genet. 44, 483–489 (2012).

11. Chatterjee, N. et al. Projecting the performance of risk prediction based on polygenic analyses of genome-wide association studies. Nat. Genet. 45, 400–5, 405e1–3 (2013).

12. Khera, A. V. et al. Genome-wide polygenic scores for common diseases identify individuals with risk equivalent to monogenic mutations. Nat. Genet. 50, 1219–1224 (2018).

13. Schumacher, F. R. et al. Association analyses of more than 140,000 men identify 63 new prostate cancer susceptibility loci. Nat. Genet. 50, 928–936 (2018).

14. Sharp, S. A. et al. Development and Standardization of an Improved Type 1 Diabetes Genetic Risk Score for Use in Newborn Screening and Incident Diagnosis. Diabetes Care 42, 200–207 (2019).

15. Kullo, I. J. et al. Incorporating a Genetic Risk Score Into Coronary Heart Disease Risk Estimates: Effect on Low-Density Lipoprotein Cholesterol Levels (the MI-GENES Clinical Trial). Circulation 133, 1181–1188 (2016).

16. Natarajan, P. et al. Polygenic Risk Score Identifies Subgroup With Higher Burden of Atherosclerosis and Greater Relative Benefit From Statin Therapy in the Primary Prevention Setting. Circulation 135, 2091–2101 (2017).

17. Márquez-Luna, C., Loh, P.-R., South Asian Type 2 Diabetes (SAT2D) Consortium, SIGMA Type 2 Diabetes Consortium & Price, A. L. Multiethnic polygenic risk scores improve risk prediction in diverse populations. Genet. Epidemiol. 41, 811–823 (2017).

18. Duncan, L. et al. Analysis of polygenic risk score usage and performance in diverse human populations. Nat. Commun. 10, 3328 (2019).

19. Hu, Y. et al. Leveraging functional annotations in genetic risk prediction for human complex diseases. PLoS Comput. Biol. 13, e1005589 (2017).

20. Márquez-Luna, C. et al. Modeling functional enrichment improves polygenic prediction accuracy in UK Biobank and 23andMe data sets. bioRxiv 375337 (2018). doi:10.1101/375337

21. Okada, Y. et al. Genetics of rheumatoid arthritis contributes to biology and drug discovery. Nature 506, 376–381 (2014).

22. Kanai, M. et al. Genetic analysis of quantitative traits in the Japanese population links cell types to complex human diseases. Nature Genetics 50, 390–400 (2018).

23. Yengo, L. et al. Meta-analysis of genome-wide association studies for height and body mass index in -700000 individuals of European ancestry. Hum. Mol. Genet. 27, 3641–3649 (2018).

24. Schaub, M. A., Boyle, A. P., Kundaje, A., Batzoglou, S. & Snyder, M. Linking disease associations with regulatory information in the human genome. Genome Res. 22, 1748–1759 (2012).

25. Maurano, M. T. et al. Systematic localization of common disease-associated variation in regulatory DNA. Science 337, 1190–1195 (2012).

26. Reshef, Y. A. et al. Detecting genome-wide directional effects of transcription factor binding on polygenic disease risk. Nat. Genet. 50, 1483–1493 (2018).

27. Liu, X., Li, Y. I. & Pritchard, J. K. Trans Effects on Gene Expression Can Drive Omnigenic Inheritance. Cell 177, 1022–1034.e6 (2019).

28. Lambert, S. A. et al. The Human Transcription Factors. Cell 172, 650–665 (2018).

29. Teytelman, L., Thurtle, D. M., Rine, J. & van Oudenaarden, A. Highly expressed loci are vulnerable to misleading ChIP localization of multiple unrelated proteins. Proc. Natl. Acad. Sci. U. S. A. 110, 18602–18607 (2013).

30. Skene, P. J. & Henikoff, S. An efficient targeted nuclease strategy for high-resolution mapping of DNA binding sites. Elife 6, (2017).

31. Amariuta, T. et al. IMPACT: Genomic Annotation of Cell-State-Specific Regulatory Elements Inferred from the Epigenome of Bound Transcription Factors. Am. J. Hum. Genet. 104, 879–895 (2019).

32. Kawakami, E., Nakaoka, S., Ohta, T. & Kitano, H. Weighted enrichment method for prediction of transcription regulators from transcriptome and global chromatin immunoprecipitation data. Nucleic Acids Res. 44, 5010–5021 (2016).

33. ENCODE Project Consortium. An integrated encyclopedia of DNA elements in the human genome. Nature 489, 57–74 (2012).

34. Roadmap Epigenomics, C. et al. Heravi-428 Moussavi A, Kheradpour P, Zhang Z, Wang J, et al. Integrative analysis of 111 reference human 429 epigenomes. Nature 518, 317–330 (2015).

35. Finucane, H. K. et al. Heritability enrichment of specifically expressed genes identifies disease-relevant tissues and cell types. Nat. Genet. 50, 621–629 (2018).

36. Gazal, S. et al. Linkage disequilibrium–dependent architecture of human complex traits shows action of negative selection. Nat. Genet. 49, 1421–1427 (2017).

37. Buniello, A. et al. The NHGRI-EBI GWAS Catalog of published genome-wide association studies, targeted arrays and summary statistics 2019. Nucleic Acids Res. 47, D1005–D1012 (2019).

38. Akiyama, M. et al. Characterizing rare and low-frequency height-associated variants in the Japanese population. Nat. Commun. 10, 4393 (2019).

39. Ishigaki, K., Akiyama, M., Kanai, M. & Takahashi, A. Large scale genome-wide association study in a Japanese population identified 45 novel susceptibility loci for 22 diseases. bioRxiv (2019).

40. Akiyama, M. et al. Genome-wide association study identifies 112 new loci for body mass index in the Japanese population. Nat. Genet. 49, 1458–1467 (2017).

41. Brown, B. C., Ye, C. J., Price, A. L. & Zaitlen, N. Transethnic Genetic-Correlation Estimates from Summary Statistics. Am. J. Hum. Genet. 99, 76–88 (2016).

42. Drake, L. Y. et al. B cells play key roles in th2-type airway immune responses in mice exposed to natural airborne allergens. PLoS One 10, e0121660 (2015).

43. Amariuta, T., Luo, Y., Knevel, R., Okada, Y. & Raychaudhuri, S. Advances in genetics toward identifying pathogenic cell states of rheumatoid arthritis. Immunol. Rev. (2019). doi:10.1111/imr.12827

44. Buttari, B., Profumo, E. & Riganò, R. Crosstalk between red blood cells and the immune system and its impact on atherosclerosis. Biomed Res. Int. 2015, 616834 (2015).

45. Anderson, H. L., Brodsky, I. E. & Mangalmurti, N. S. The Evolving Erythrocyte: Red Blood Cells as Modulators of Innate Immunity. J. Immunol. 201, 1343–1351 (2018).

46. Lui, J. C. & Baron, J. Mechanisms limiting body growth in mammals. Endocr. Rev. 32, 422–440 (2011).

47. Maier, A. B., van Heemst, D. & Westendorp, R. G. J. Relation between body height and replicative capacity of human fibroblasts in nonagenarians. J. Gerontol. A Biol. Sci. Med. Sci. 63, 43–45 (2008).

48. Murphy, R. A. et al. Adipose tissue, muscle, and function: potential mediators of associations between body weight and mortality in older adults with type 2 diabetes. Diabetes Care 37, 3213–3219 (2014).

49. Heymsfield, S. B., Gallagher, D., Mayer, L., Beetsch, J. & Pietrobelli, A. Scaling of human body composition to stature: new insights into body mass index. Am. J. Clin. Nutr. 86, 82–91 (2007).

50. Kichaev, G. et al. Leveraging Polygenic Functional Enrichment to Improve GWAS Power. Am. J. Hum. Genet. 104, 65–75 (2019).

51. Gusev, A. et al. Atlas of prostate cancer heritability in European and African-American men pinpoints tissue-specific regulation. Nat. Commun. 7, 10979 (2016).

52. Gibbs, R. A. et al. A global reference for human genetic variation. Nature 526, 68–74 (2015).

53. Nagai, A. et al. Overview of the BioBank Japan Project: Study design and profile. J. Epidemiol. 27, S2–S8 (2017).

54. Hirata, M. et al. Cross-sectional analysis of BioBank Japan clinical data: A large cohort of 200,000 patients with 47 common diseases. J. Epidemiol. 27, S9–S21 (2017).

## Online Methods References

1. Kawakami, E., Nakaoka, S., Ohta, T. & Kitano, H. Weighted enrichment method for prediction of transcription regulators from transcriptome and global chromatin immunoprecipitation data. Nucleic Acids Res. 44, 5010–5021 (2016).

2. Ishigaki, K., Akiyama, M., Kanai, M. & Takahashi, A. Large scale genome-wide association study in a Japanese population identified 45 novel susceptibility loci for 22 diseases. bioRxiv (2019).

3. Amariuta, T. et al. IMPACT: Genomic Annotation of Cell-State-Specific Regulatory Elements Inferred from the Epigenome of Bound Transcription Factors. Am. J. Hum. Genet. 104, 879–895 (2019).

4. ENCODE Project Consortium. An integrated encyclopedia of DNA elements in the human genome. Nature 489, 57–74 (2012).

5. Kanai, M. et al. Genetic analysis of quantitative traits in the Japanese population links cell types to complex human diseases. Nat. Genet. 50, 390–400 (2018).

6. Gazal, S. et al. Linkage disequilibrium–dependent architecture of human complex traits shows action of negative selection. Nat. Genet. 49, 1421–1427 (2017).

7. Finucane, H. K. et al. Partitioning heritability by functional annotation using genome-wide association summary statistics. Nat. Genet. 47, 1228–1235 (2015).

8. Finucane, H. K. et al. Heritability enrichment of specifically expressed genes identifies disease-relevant tissues and cell types. Nat. Genet. 50, 621–629 (2018).

9. Gibbs, R. A. et al. A global reference for human genetic variation. Nature 526, 68–74 (2015).

10. Nagai, A. et al. Overview of the BioBank Japan Project: Study design and profile. J. Epidemiol. 27, S2–S8 (2017).

11. Hirata, M. et al. Cross-sectional analysis of BioBank Japan clinical data: A large cohort of 200,000 patients with 47 common diseases. J. Epidemiol. 27, S9–S21 (2017).

12. Heinz, S. et al. Simple combinations of lineage-determining transcription factors prime cis-regulatory elements required for macrophage and B cell identities. Mol. Cell 38, 576–589 (2010).

13. Brown, B. C., Ye, C. J., Price, A. L., Zaitlen, N. & Asian Genetic Epidemiology Network-Type 2 Diabetes. Transethnic genetic correlation estimates from summary statistics. doi:10.1101/036657

14. Martin, A. R. et al. Clinical use of current polygenic risk scores may exacerbate health disparities. Nat. Genet. 51, 584–591 (2019).

15. Purcell, S. et al. PLINK: a tool set for whole-genome association and population-based linkage analyses. Am. J. Hum. Genet. 81, 559–575 (2007).

16. Lam, M. et al. Comparative genetic architectures of schizophrenia in East Asian and European populations. doi:10.1101/445874

17. Lee, S. H., Goddard, M. E., Wray, N. R. & Visscher, P. M. A better coefficient of determination for genetic profile analysis. Genet. Epidemiol. 36, 214–224 (2012).

