## Supplemental Text and Figures for "*In silico* integration of thousands of epigenetic datasets into 707 cell type regulatory annotations improves the trans-ethnic portability of polygenic risk scores"

### Supplementary Figures

### Figure S1

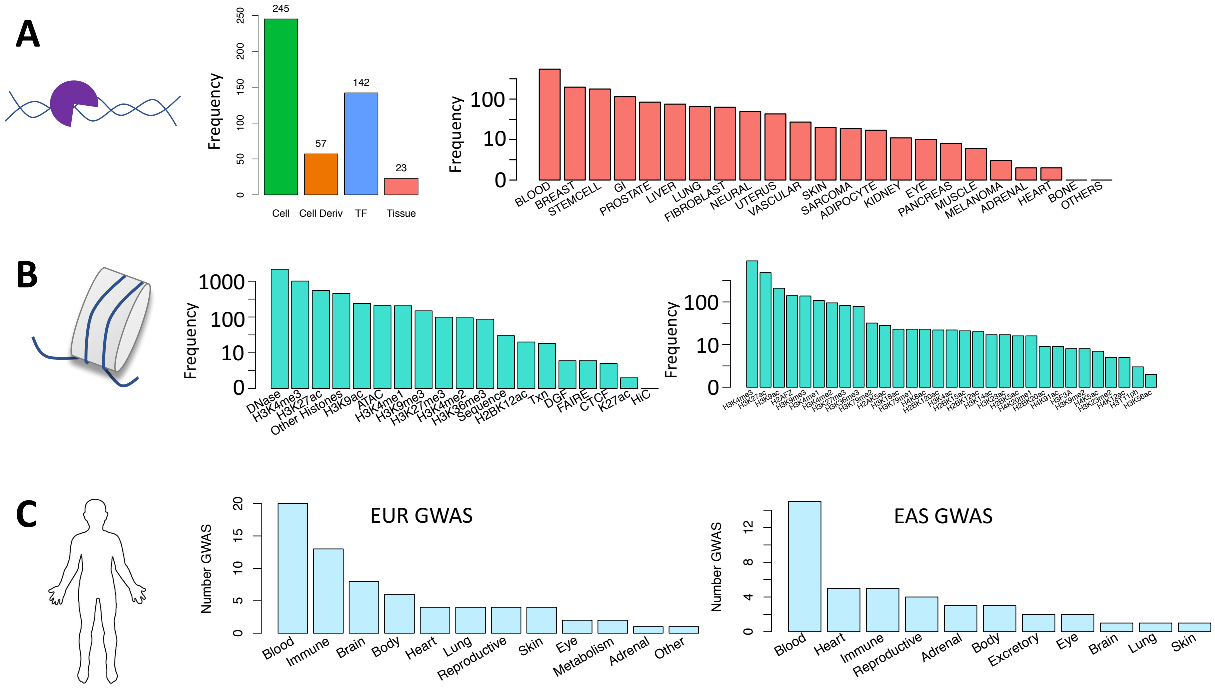


Figure S1 legend. A) TF ChIP-seq collection from NCBI: (left) cell type and TF diversity where "Cell Deriv" indicates number of unique parental cell types, for example GM12878 and GM10847 are both B cell lines, (right) diversity of tissue types. B) (left) Epigenomic and sequence features to be used in IMPACT model, (right) diversity of histone modification ChIP-seq in features. C) Diversity of European (EUR) and East Asian (EAS) GWAS summary statistics across phenotypic categories.

### Figure S2

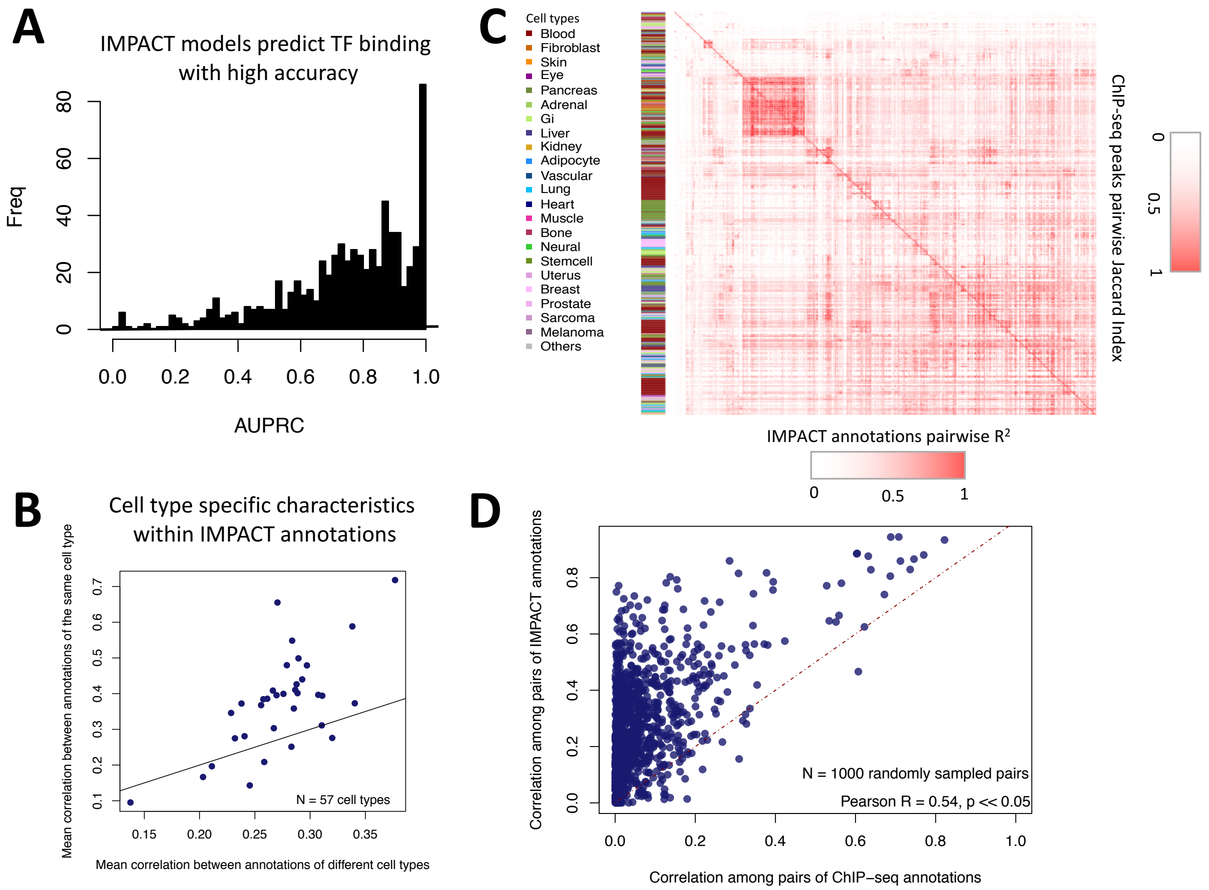


Figure S2 legend. A) Histogram of prediction performance of 707 IMPACT models (metric = AUPRC). B) IMPACT annotations of the same cell type are more similar to one another than annotations of different cell types. C) Pairwise correlation of IMPACT regulatory element annotations (lower triangle of matrix) relative to pairwise correlation of corresponding TF ChIP-seq annotations (upper triangle of matrix). Pearson r was calculated using probabilities assigned to 779,355 SNPs on chr1 from phase 3 of 1000G (EUR), Jaccard indices were calculated for binary ChIP-seq tracks genome-wide, in which the size of the intersection of base pairs between two datasets was divided by the size of the union of base pairs. D) Pairwise correlations between 1000 randomly selected datasets between TF ChIP-seq and their corresponding IMPACT annotations; values sampled from C).

**Figure S3**

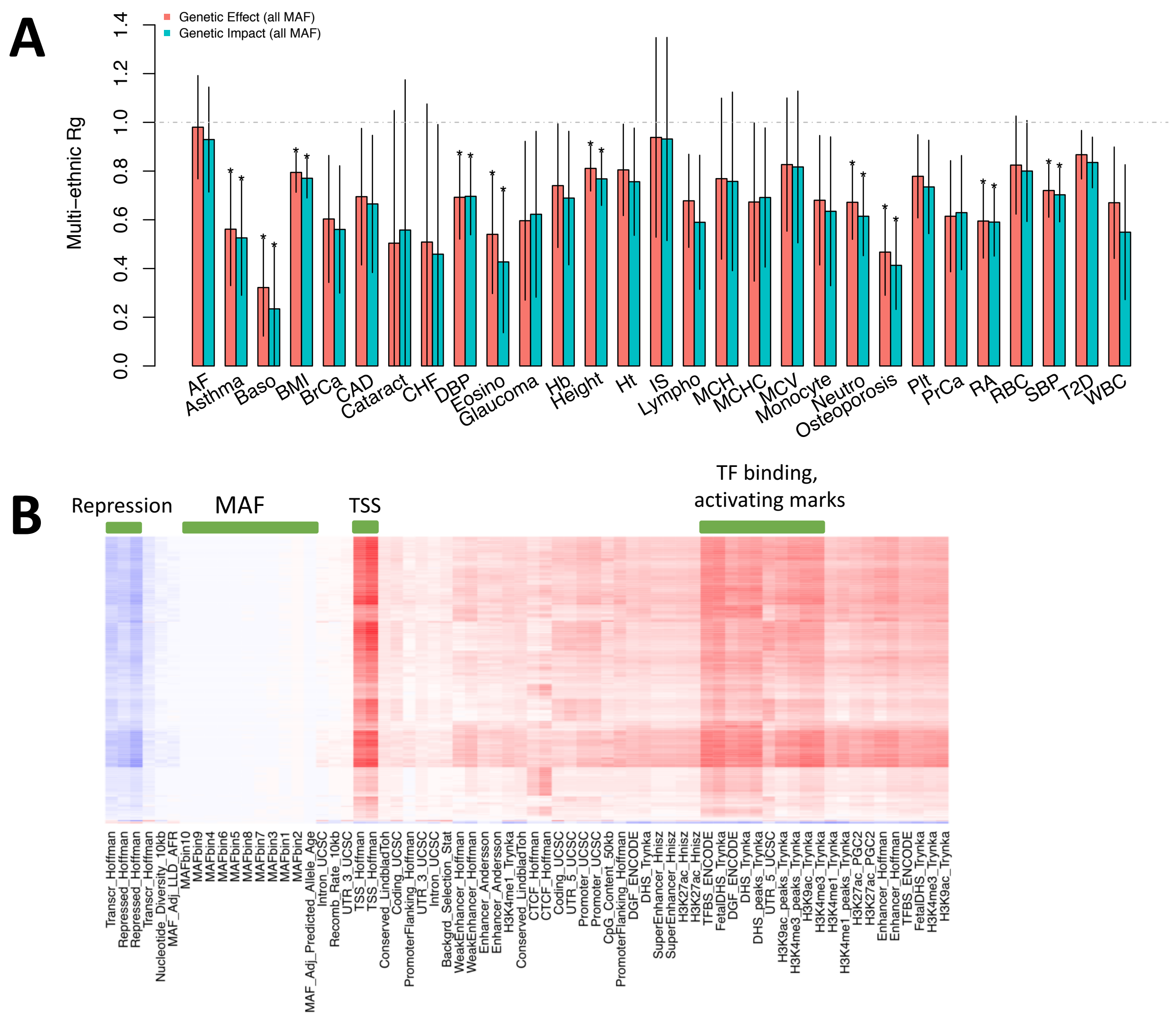


Figure S3 legend. A) For the 29 traits for which we collected both EUR and EAS GWAS summary statistics, we computed the multi-ethnic genetic correlation with Popcorn. For 10 traits, the genetic correlation is significantly less than 1, indicated by an asterisk (*P* < 0.05 / 29 traits). We plot both the genetic correlation computed separately using genetic effect (effect size estimates not normalized to allele frequency) and genetic impact (allele variance normalized effect sizes). B) IMPACT annotations correlate most with TSS, TFBS, and activation histone mark annotations, while no correlation is present with European ancestry MAF bins.

**Figure S4**

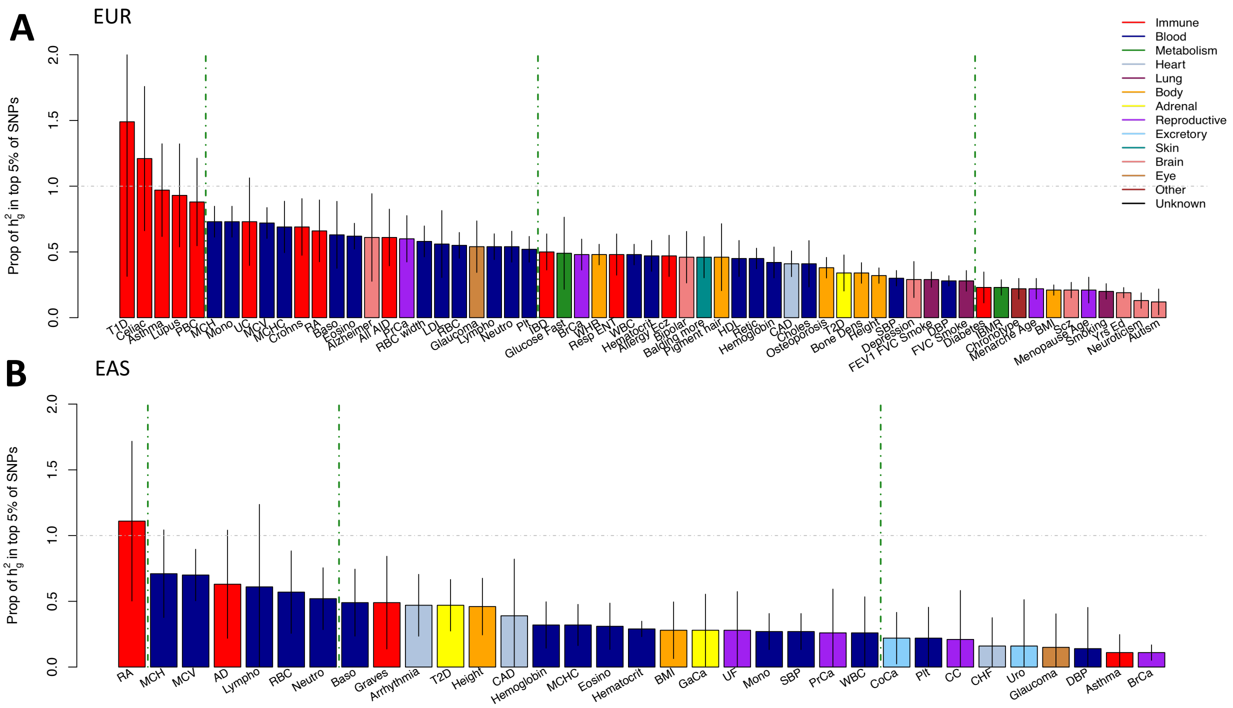


Figure S4 legend. A) Common SNP heritability captured by the top 5% of SNPs according to the lead cell type association for each EUR GWAS. Lead association determined by largest $\tau^*$ estimate that is significantly positive. B) Similar for each EAS GWAS. Gray bars indicate the standard error of the heritability estimate. Color represents the category of the complex trait or disease.

**Figure S5**

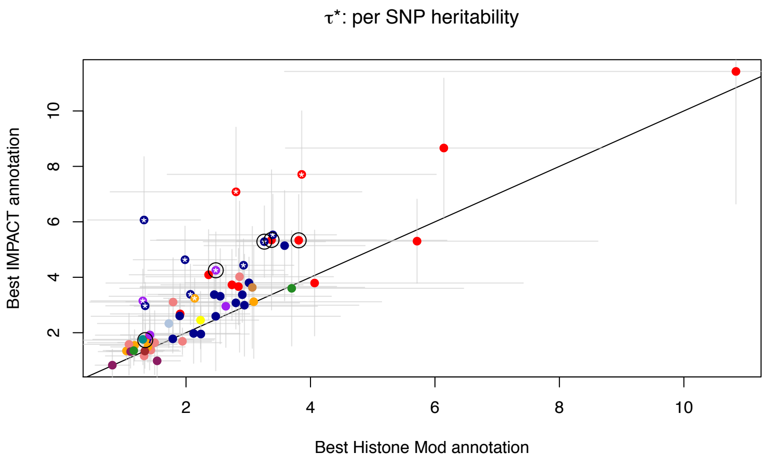


Figure S5 legend. Comparison of two different functional annotations, IMPACT and cell type specific histone marks, to capture polygenic heritability assessed by quantifying $\tau^*$ per-SNP heritability value. Circled are five representative traits used throughout the study: asthma, RA, PrCa, MCV, and height.

**Figure S6**

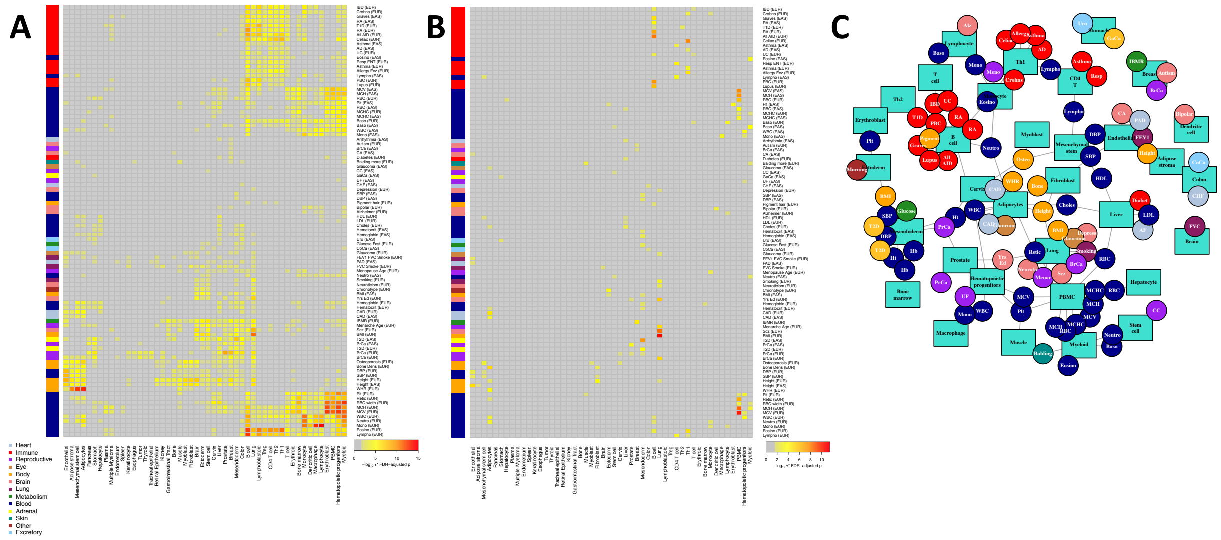


Figure S6 legend. A) Stratification of IMPACT annotation associations by 50 cell types across the 95 polygenic traits and diseases of 111 with at least one association. For each cell type, the strongest annotation association is represented (-$\log_{10}$ $\tau^*$ $P$ value, FDR 5% adjusted). B) After four rounds of conditional analysis, non-independent associations were removed. Shown are the remaining independent annotation associations of the same 50 cell types and 95 traits. Color indicates -$\log_{10}$ $\tau^*$ $P$ value adjusted for FDR 5%; if more than one independent cell type association, -$\log_{10}$ $\tau^*$ conditional $P$ value adjusted for FDR 5% is indicated. C) Network of remaining independent associations, same information as in B), reveals clusters of regulatory modules that recapitulate known biology.

**Figure S7**

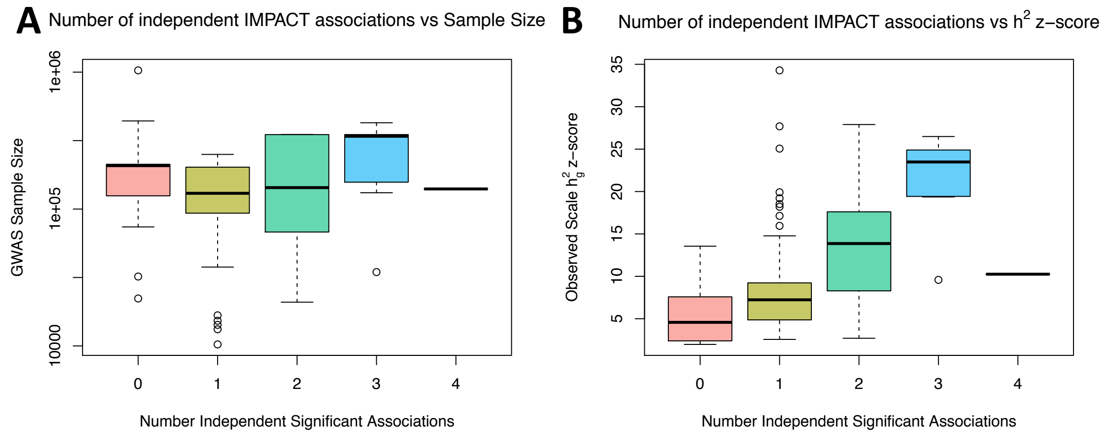


Figure S7 legend.

A) Number of independent IMPACT cell type associations is not significantly correlated with the sample size of the GWAS ($P$ = 0.19). B) Number of independent associations is significantly positively correlated with the observed scale heritability z-score of the trait ($P$ < 5.4e-9).

**Figure S8**

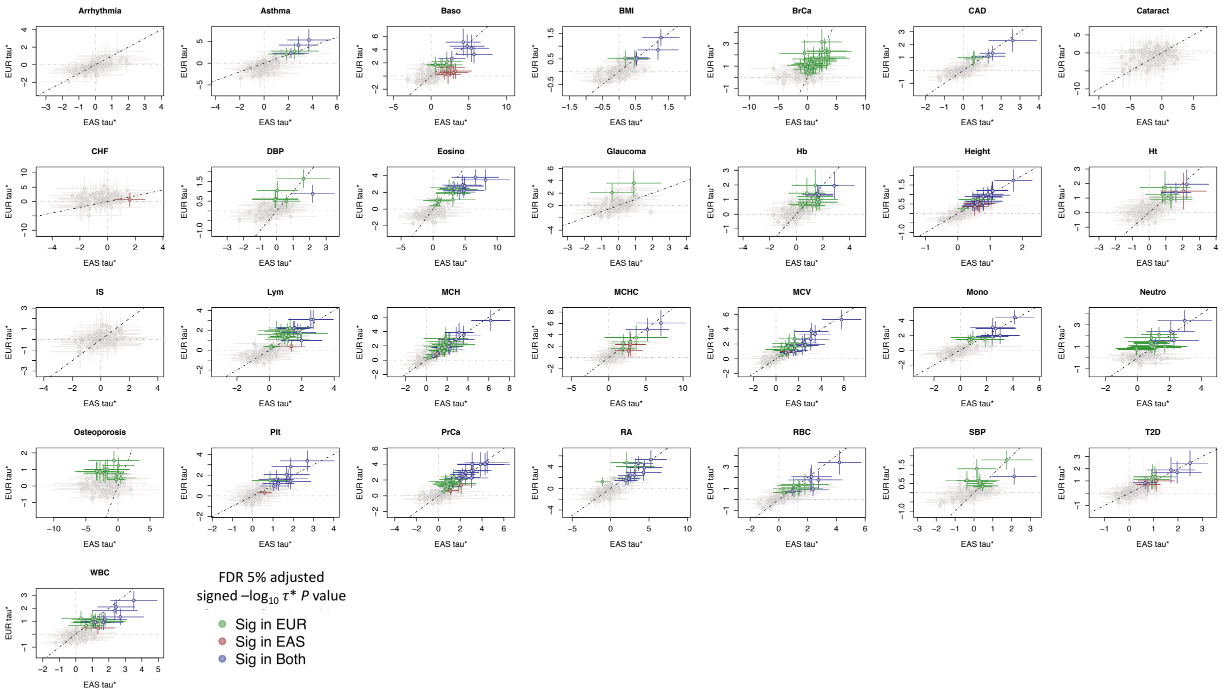


Figure S8 legend. Common per-SNP heritability ($\tau^*$) estimate for sets of independent IMPACT cell type annotations across 29 traits. Dotted line is the identity line, y = x. $\tau^*$ values with their standard errors are colored green if significantly positive in EUR and not EAS, red if significantly positive in EAS but not in EUR, green if significantly positive in both EUR and EAS, and gray if not significantly positive in either population.

**Figure S9**

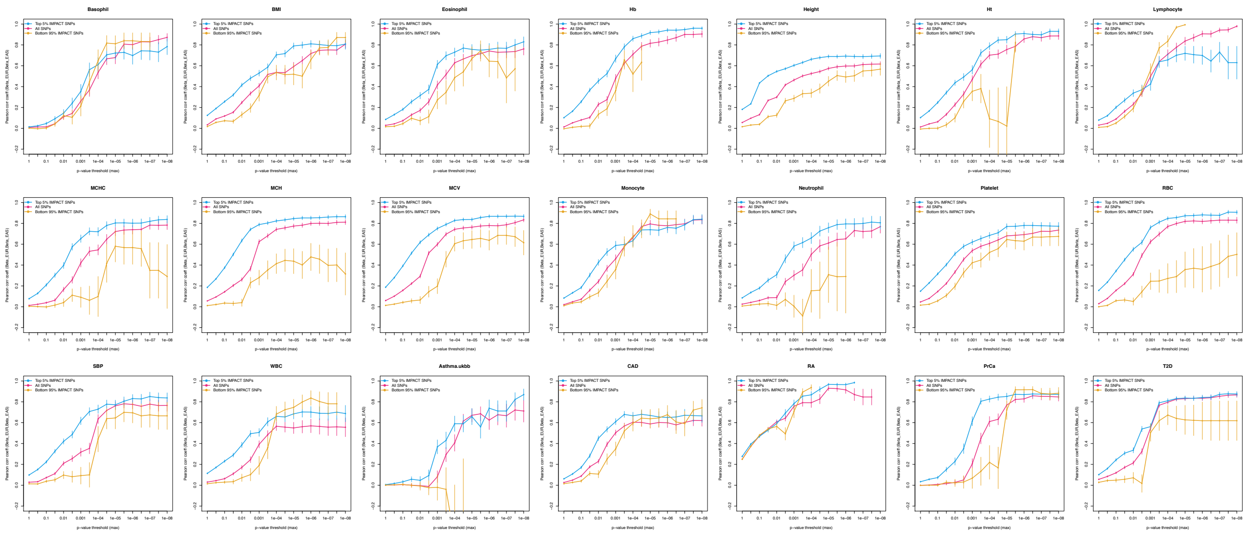


Figure S9 legend. For 21 traits shared between EUR and EAS, effect size correlation (Pearson correlation coefficient) across 17 *P* value thresholds for three partitions of SNPs genome-wide: 1) lead SNPs with no IMPACT inference (yellow), 2) top 5% of SNPs according to the largest $\tau^*$ effect size IMPACT annotation (red), and 3) the bottom 95% of SNPs according to the same IMPACT annotation (teal). Vertical lines indicate one standard deviation of the correlation coefficient estimate.

**Figure S10**

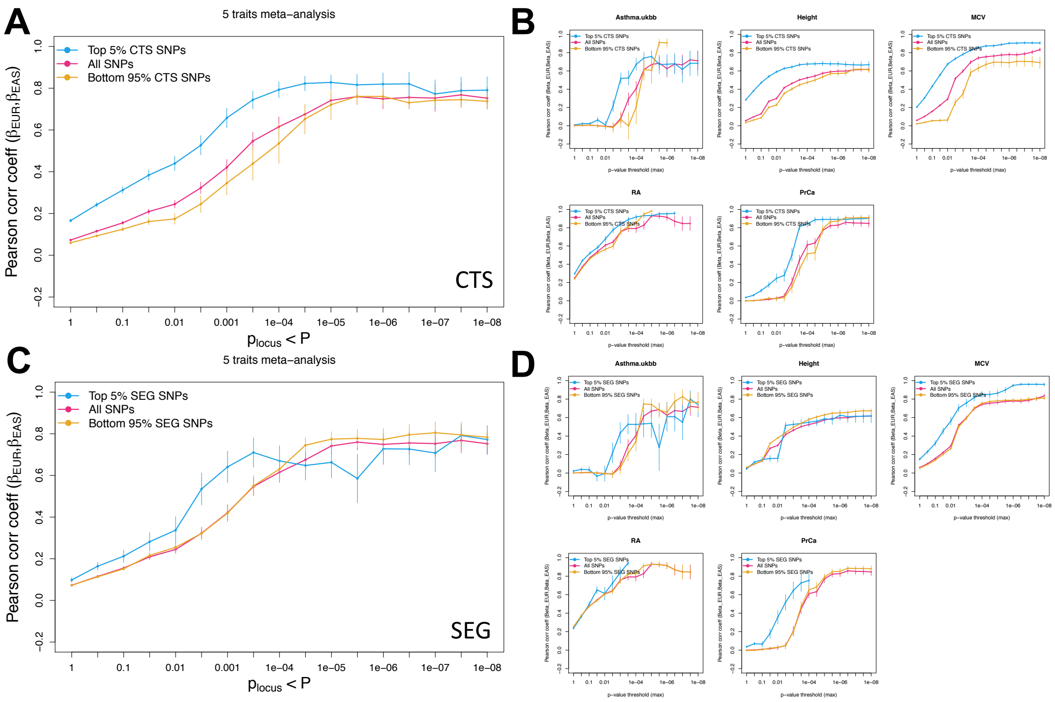


Figure S10 legend. For 5 traits representing different biological underpinnings shared between EUR and EAS (subset of 21 investigated in our study), we report the effect size correlation (Pearson correlation coefficient) across 17 *P* value thresholds for three partitions of SNPs genome-wide: 1) lead SNPs with no functional inference (yellow), 2) top 5% of SNPs according to the largest $\tau^*$ annotation effect size (red), and 3) the bottom 95% of SNPs according to the same functional annotations (teal). Here, we select the top annotation in two categories of previously published functional annotations: first, from LDSC CTS annotations (meta-analysis in A, individual traits in B) and second, from LDSC SEG annotations (meta-analysis in C, individual traits in D). Vertical lines indicate two standard deviations of the correlation coefficient estimate.

**Figure S11**

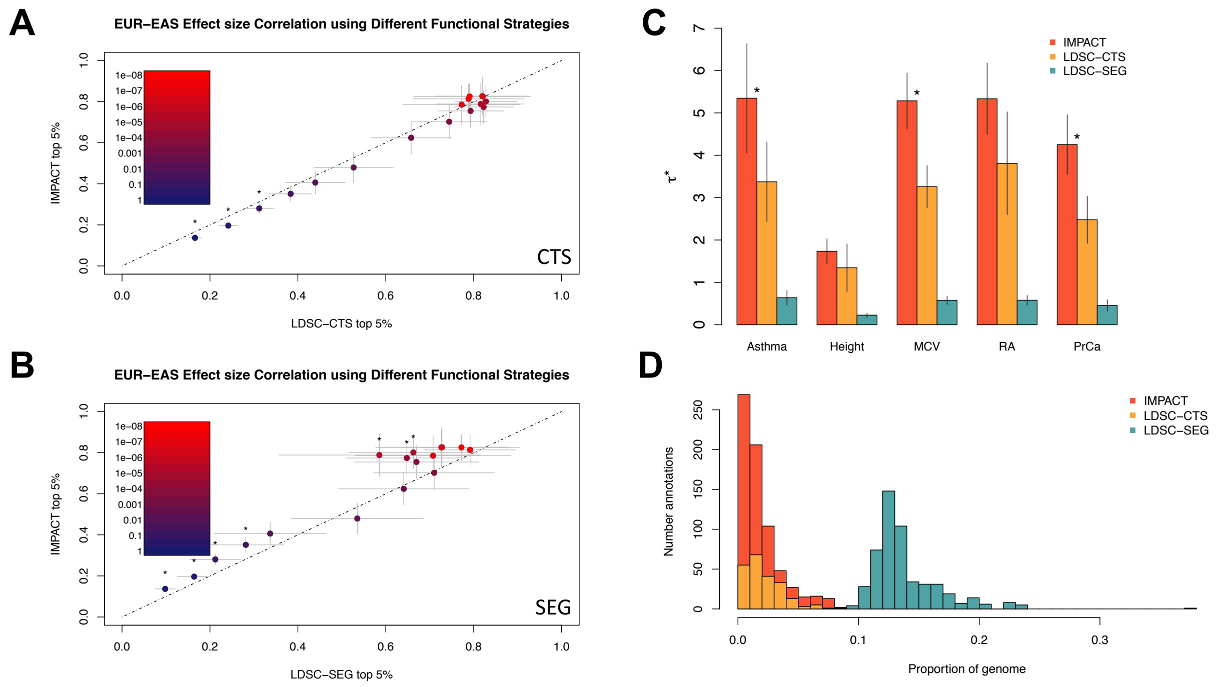


Figure S11 legend. A) Comparison of top LDSC-CTS annotations in multi-ethnic effect size correlation analysis with top IMPACT annotations meta-analyzed over 5 traits. Asterisks indicate three p-value thresholds at which top LDSC-CTS annotations outperformed top IMPACT annotations (n = 3, difference of means test at $P < 0.05$). B) Similar to A) but for LDSC-SEG annotations. Asterisks indicate three p-value thresholds at which IMPACT annotations outperformed top LDSC-SEG annotations (n = 13, difference of means test at p < 0.05). C) $\tau^*$ common per-SNP heritability captured genome-wide across the 5 selected traits reveals that IMPACT annotations generally capture more heritability than LDSC-CTS annotations (indicated by asterisk, difference of means test $P < 0.05$) and consistently more than LDSC-SEG annotations. D) Distribution of annotation sizes for three different functional regimes: IMPACT (red), LDSC-CTS (yellow), LDSC-SEG (teal).

**Figure S12**

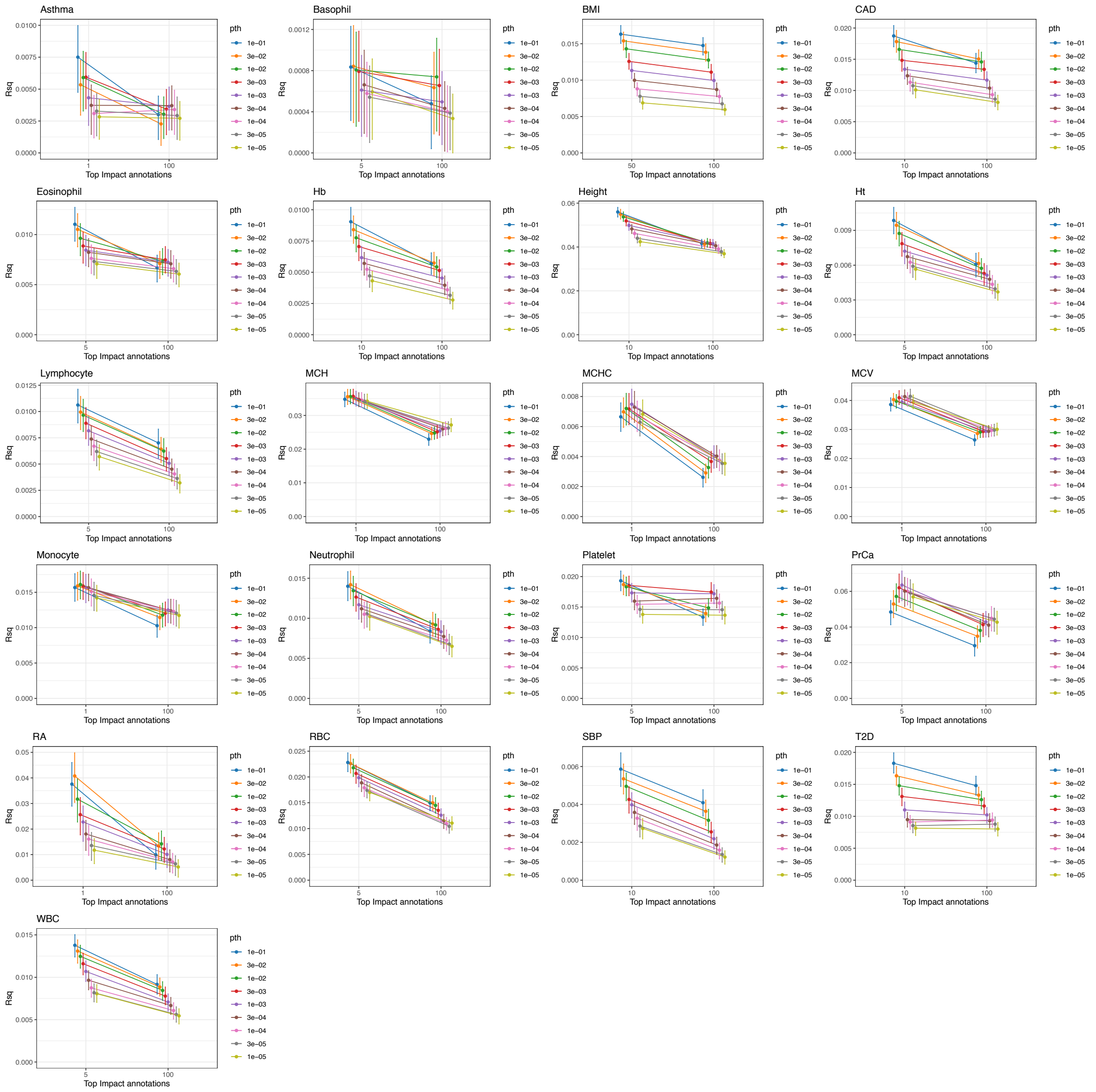


Figure S12 legend. EUR PRS model evaluated on EAS individuals from BBJ. For each trait, we evaluate the predictive value of standard PRS models (top 100% of IMPACT SNPs) and functionally-informed PRS models (using a subset of SNPs prioritized by IMPACT). Intervals represent the 95% confidence interval around the $R^2$ estimate. For quantitative traits, $R^2$ represents the proportion of variance captured by the linear PRS model. For case control traits, $R^2$ represents the liability scale $R^2$ from the logistic regression PRS model.

**Figure S13**

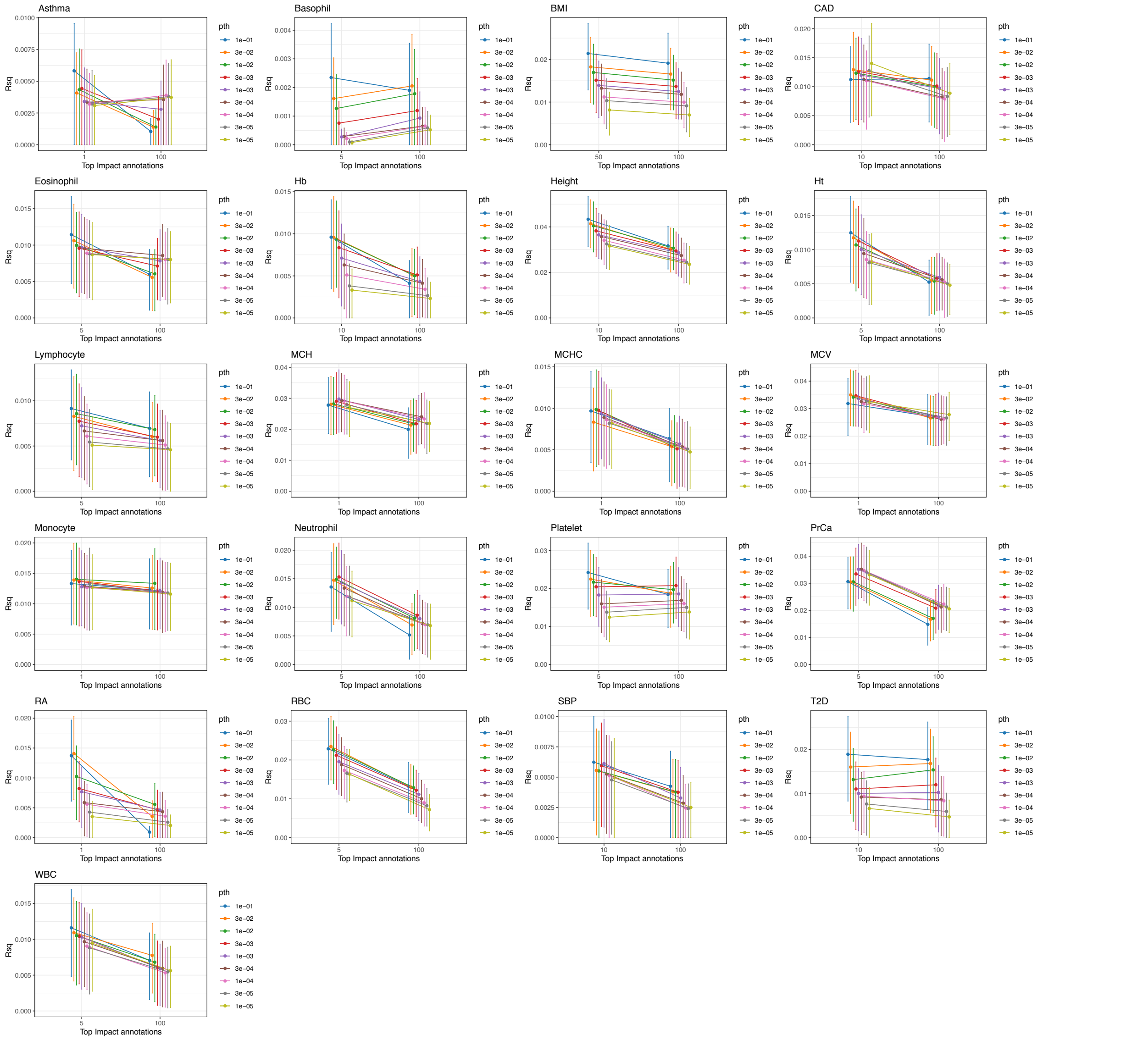


Figure S13 legend. EUR PRS model evaluated on 5,000 randomly selected EAS individuals from BBJ. For each trait, we evaluate the predictive value of standard PRS models (top 100% of IMPACT SNPs) and functionally-informed PRS models (using a subset of SNPs prioritized by IMPACT). Intervals represent the 95% confidence interval around the $R^2$ estimate. For quantitative traits, $R^2$ represents the proportion of variance captured by the linear PRS model.

For case control traits, $R^2$ represents the liability scale $R^2$ from the logistic regression PRS model.

**Figure S14**

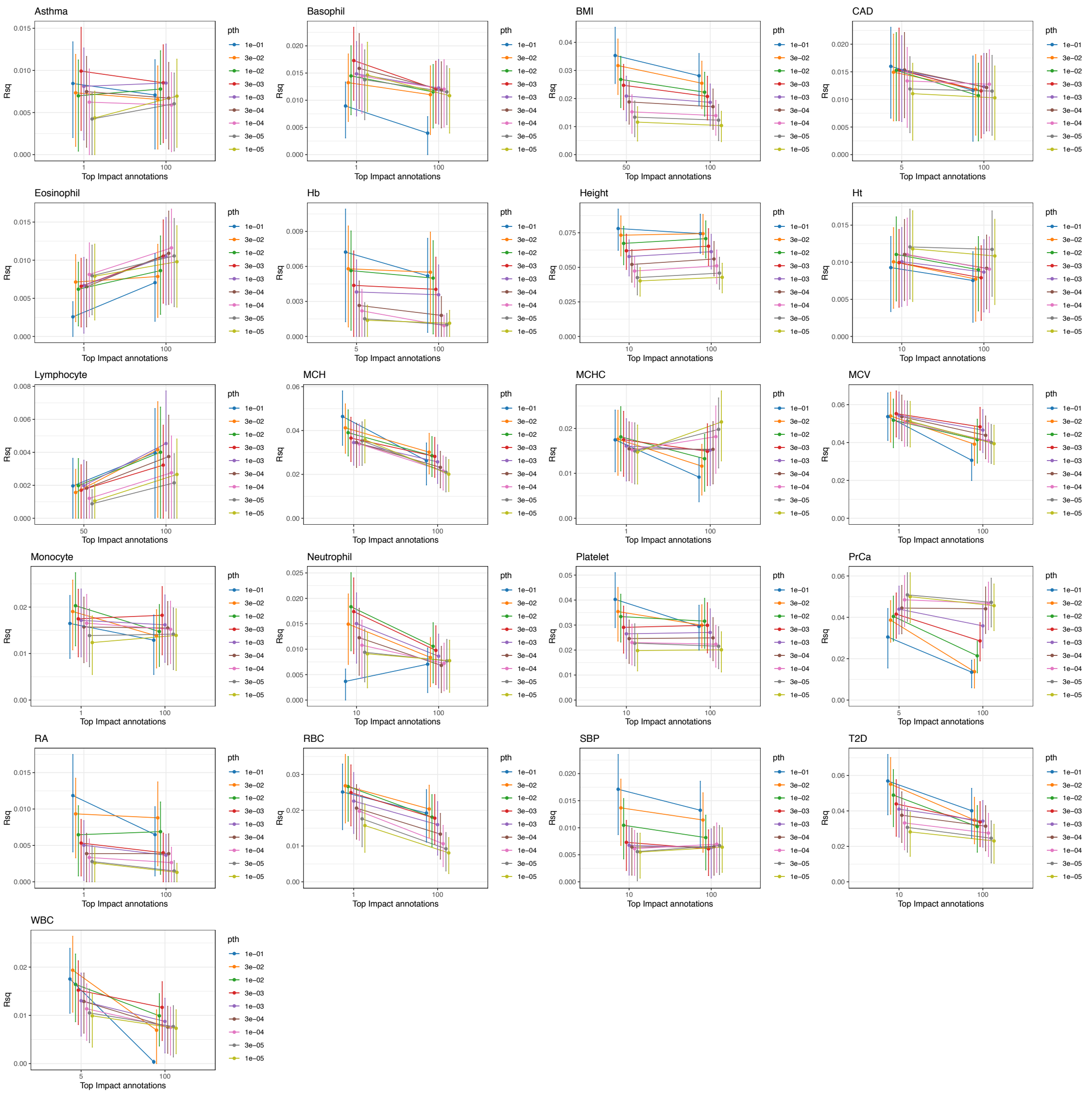


Figure S14 legend. EAS PRS model evaluated on 5,000 non-overlapping EAS individuals from BBJ; these 5,000 individuals are the same as EAS test individuals in SF13 and SF15. For each trait, we evaluate the predictive value of standard PRS models (top 100% of IMPACT SNPs) and functionally-informed PRS models (using a subset of SNPs prioritized by IMPACT). Intervals represent the 95% confidence interval around the $R^2$ estimate. For quantitative traits, $R^2$ represents the proportion of variance captured by the linear PRS model. For case control traits, $R^2$ represents the liability scale $R^2$ from the logistic regression PRS model.

**Figure S15**

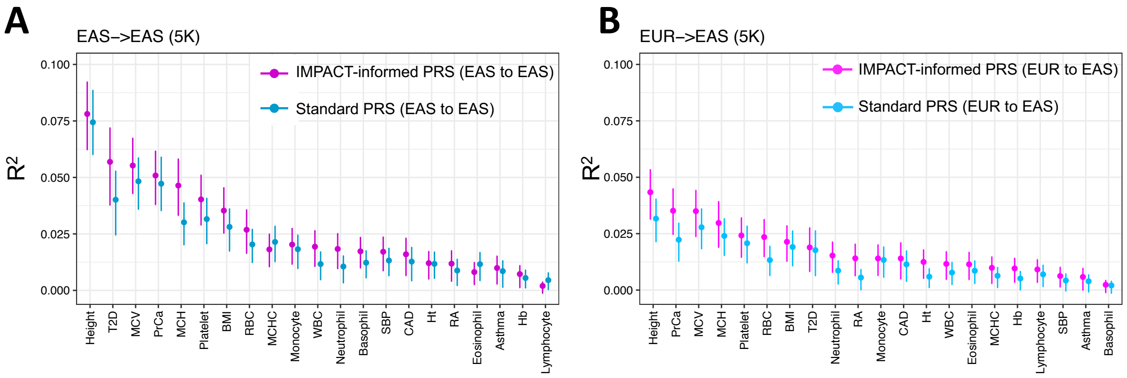


Figure S15 legend. A) Phenotypic variance ($R^2$) in 5,000 BBJ individuals explained by IMPACT-informed PRS-EAS (dark pink) and standard PRS-EAS (dark blue). B) Phenotypic variance ($R^2$) in 5,000 BBJ individuals explained by IMPACT-informed PRS-EUR (light pink) and standard PRS-EUR (light blue). Error bars indicate 95% CI calculated via 1,000 bootstraps.

**Figure S16**

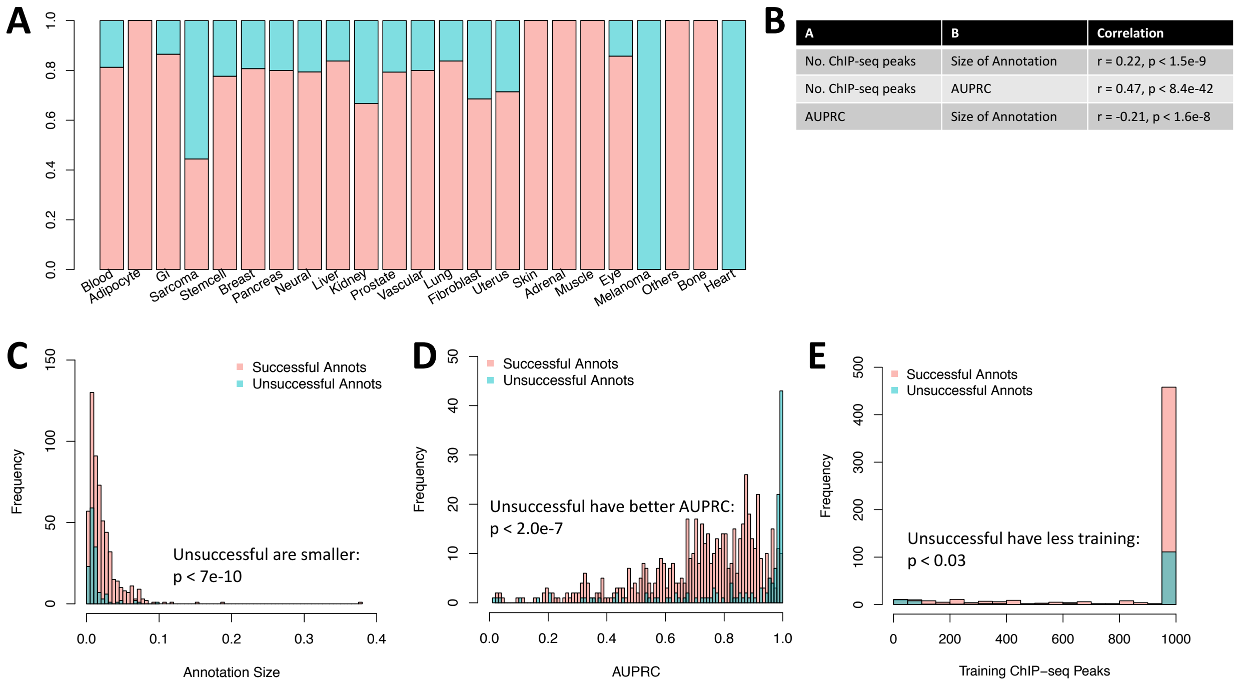


Figure S16 legend. A) Distribution of annotation size (average IMPACT score over annotated SNPs) for "successful" and "unsuccessful" annotations. B) Distribution of TF binding model AUPRC for "successful" and "unsuccessful" annotations. C) Distribution of training set size (number of TF ChIP-seq peaks) for "successful" and "unsuccessful" annotations. D) Correlation of metadata factors of IMPACT annotations: number of ChIP-seq peaks available to training data, AUPRC of TF binding prediction model, and annotation size. E) For each tissue type category of IMPACT annotation, the proportion of annotations that were significantly associated with at least one polygenic trait or disease ("successful") is indicated by the height of the pink bar. "Unsuccessful" annotations were not found to be significantly associated with any phenotype and are indicated by the green bar. For example, heart-labeled annotations had no significant associations.

**Figure S17**

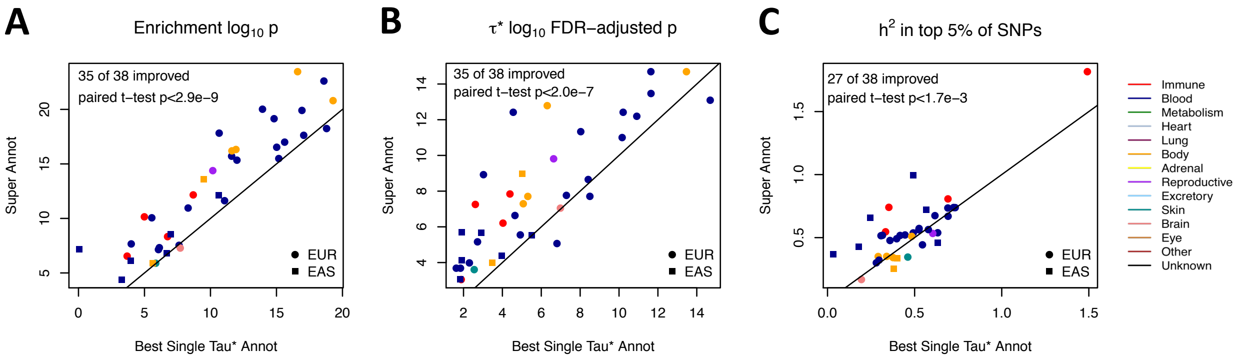


Figure S17 legend. Comparison of heritability metrics between the lead annotation and the composite annotation, created from independently associated IMPACT annotations. A) Statistical significance of the enrichment estimate. B) Statistical significance of the $\tau^*$ S-LDSC regression coefficient estimate. C) Proportion of observed scaled heritability in the top 5% SNPs scored by IMPACT.

**Figure 18**

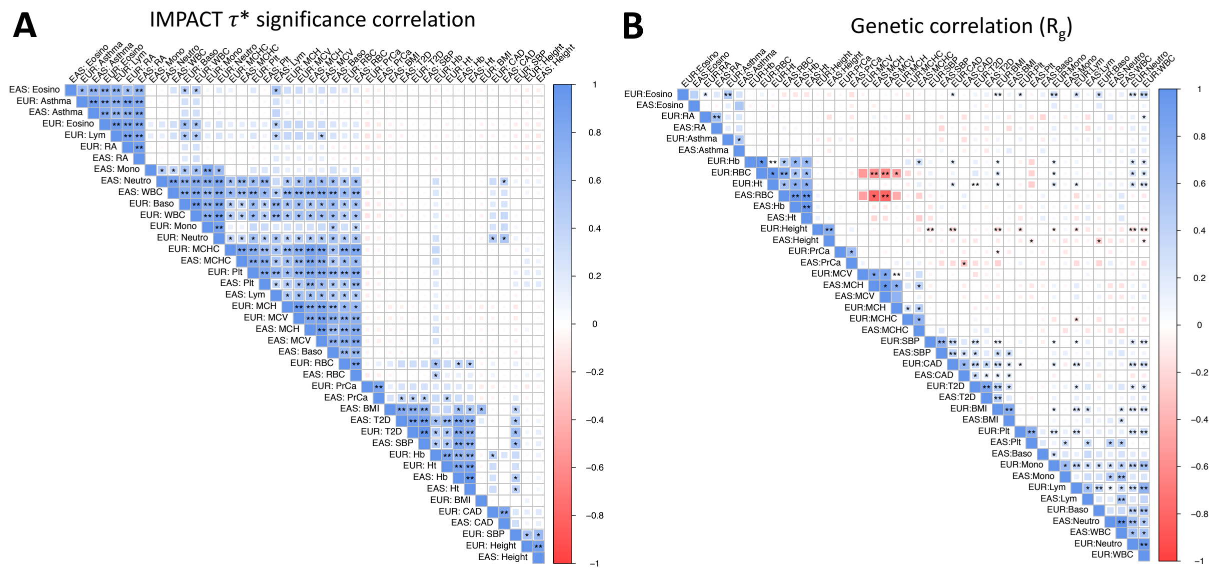

Figure S18 legend. A) Pairwise correlation of IMPACT functional annotations' $\tau^*$ significance across 42 traits, accounting for 21 unique phenotypes (those with at least one significant IMPACT association in both EUR and EAS) and two populations. * indicates FDR-adjusted *P* < 0.05, ** indicates FDR-adjusted *P* < 1e-10. B) Pairwise genetic correlation across the same 42 traits as in (A). * indicates nominal *P* < 0.05, ** indicates nominal *P* < 1e-10.

**Figure S19**

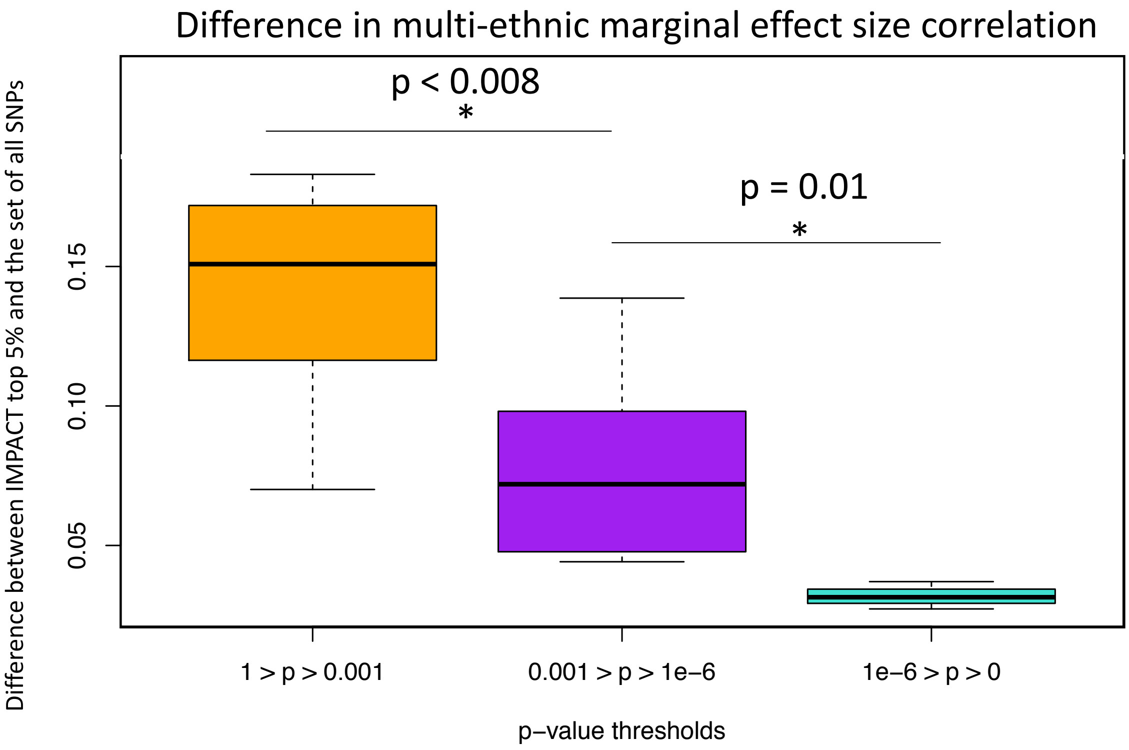


Figure S19 legend. Improvement by functional data (IMPACT top 5% SNP selection) varies by *P* value threshold. Improvement is greatest when p-values are lenient (orange). Improvement is minimized when the EUR GWAS *P* value is near or past the genome-wide significant threshold (turquoise).

**Extended Data Figure 1**

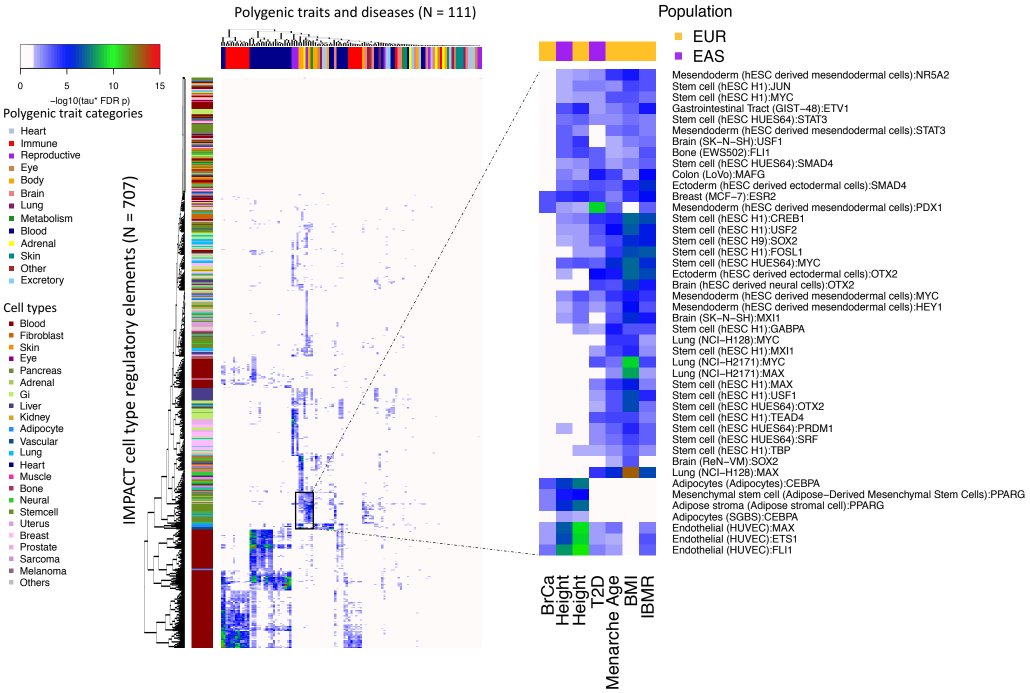


Ext. Data 1 legend. Significant cell type-phenotype associations across 707 IMPACT regulatory annotations and 111 complex traits and diseases at $\tau^*$ 5% FDR, color indicates -log10 FDR 5% adjusted *P* value of $\tau^*$. Zooms shows particular cell type categories enriched for polygenic trait associations.

**Supplemental Text**

<u>Significant IMPACT annotation-trait associations</u>

We identified at least one statistically significant IMPACT annotation association with 95 of 111 polygenic traits. These 95 account for 60 of 69 European phenotypes and 35 of 42 East Asian phenotypes. Analogously, across 707 cell type regulatory annotations, we identified at least one significant annotation-trait association for 566 annotations at 5% FDR. For all trait-annotation pairs, the computed $\tau^*$ and enrichment estimates, along with their standard errors can be found in **ST4-8**.

Annotations and traits with no observed heritability enrichment

For 16 polygenic traits, we observed no statistically significant annotation association. Of these 16 polygenic traits, 9 were from European GWAS; these are anorexia, cataract, “ever smoked”, three pigmentation phenotypes (skin, sunburn, tanning), and three heart disease phenotypes (CHF, IS, AF). The remaining 7 traits with no annotation associations from East Asian GWAS were cataract, COPD, IS, keloid, osteoporosis, pancreatic cancer, and pollinosis. Likewise, for 141 IMPACT annotations, we observed  no statistically significant trait association. These annotations included melanoma and heart-labeled annotations (**SF16**). Just over 40% of sarcoma annotations were significantly associated with at least one trait; for all other tissue types, more than 60% of the corresponding annotations were significantly associated with at least one trait. We found that number of training ChIP-seq peaks were significantly correlated with both the size of annotation and the AUPRC of the TF binding model (Pearson r = 0.22, $P$ < 1.5e-9; Pearson r = 0.47, $P$ < 8.4e-42, respectively) (**SF15**). However, the AUPRC and size of annotation are significantly negatively correlated (Pearson r = -0.21, $P$ < 1.6e-8). This perhaps indicates that models with a small number of training peaks and above-average AUPRC (overfitting) will lead to smaller annotations which don’t adequately cover the polygenic space, leading to fewer significant heritability enrichments. Moreover, we found that these unassociated annotations have generally  significantly smaller annotation sizes ($P$ < 7.0e-10), significantly higher TF binding model AUPRCs (*P* < 2.0e-7), significantly less training data (*P* < 0.03), and are biased for particular cell types (**SF16**).

Conditional S-LDSC analysis to identify independent annotation-trait associations

Before performing serial conditional analyses, for 9 polygenic traits, we observed a single associated cell type: EUR autism (breast), EAS breast cancer (breast), EAS cervical cancer (stem cell), EAS congestive heart failure (colon), EAS diastolic blood pressure (mesendoderm), EAS gastric cancer (stomach), EAS glaucoma (adipocytes), EAS systolic blood pressure (mesendoderm), EAS uterine fibroids (hematopoietic progenitors). However, for 86 traits, we observed that regulatory elements of multiple IMPACT annotations, mostly implicating diverse cell types, significantly capture heritability (**SF6**). After performing serial conditional analyses to resolve dependent and independent associations, there remained a total of 142 independent cell type-trait associations (**SF6**): 1 trait with 4 associations, 7 traits with 3, 30 traits with 2, 57 traits with 1, and 16 traits with none. Four annotations independently explained significant proportions of heritability in EUR prostate cancer: prostate (TFAP4), prostate (RUNX2), mesendoderm (PDX1), and cervix (NFYB). For seven European traits, three IMPACT annotations independently captured polygenic heritability: height (adipocytes, fibroblasts, lung), neutrophil count (monocytes, adipocytes, B cells), osteoporosis (myoblasts, mesenchymal stem cells, cervix), IBD (T cells and two B cell annotations), platelet count (PBMCs, hematopoietic progenitors, muscle), systolic blood pressure (endothelial, mesenchymal stem cells, fibroblasts), and white blood cell count (B cells, adipocytes, hematopoietic progenitors). For each of 22 European traits and 8 East Asian traits, we observed exactly two independent IMPACT annotation associations. Finally, for each of 30 European traits and 27 East Asian traits, we observed exactly one independent IMPACT association. For Crohn's (EUR), Th1s and naive CD4+ T cells independently captured heritability, suggesting two different biological mechanisms one via naive T cells and the other via memory effector cells. Although previous studies suggested an important role of T cells in UC[1], our study identified not only T cells but also B cells as contributors to disease pathogenesis. For UC (EUR), T cells and B cells contribute independently to explain heritability. In summary, we have elucidated the biology of some polygenic traits through resolving not only the most significantly associated cell type, but also secondary, tertiary, and quaternary independent mechanisms. These results also shed light on shared regulatory programs between cell types: in cases where prior to conditioning, we observed many diverse cell type associations, yet upon conditioning revealed a single independent signal. For example, in EUR RA, B cells were most strongly associated, while CD4+ memory T cell annotations also captured significant proportions of heritability. However, these T cell annotations were not associated independently of B cells, suggesting that RA heritability resides in shared regulatory elements between T and B cells. In summary, we have elucidated the biology of some polygenic traits through resolving not only the most significantly associated cell type, but also secondary, tertiary, and quaternary independent mechanisms.

We note that our cell type interpretations above rely on the fidelity of the IMPACT model to accurately predict TF binding in the desired cell type; a poor model may learn an epigenetic signature that does not represent the desired cell type. For identified independent cell type associations, IMPACT prediction accuracy was reasonably high: mean AUPRC = 0.67 (sd =

0.008). This distribution of AUPRC values is significantly less than the distribution of all IMPACT annotations ($P < 2.4e-3$), where the mean AUPRC is 0.74 (sd = 0.008), ranging from 0.018 to 0.998. This is consistent with our observation that IMPACT annotations with nearly perfect AUPRCs are less likely to capture polygenic heritability (**SF16**).

#### Cell type composite annotations targeting multiple independent mechanisms of polygenic traits

In light of observing 38 phenotypes for which multiple cell type regulatory element annotations independently captured significant proportions of heritability, we created composite cell type annotations in hopes of improving heritability capture. For example, we observed that genetic variation governing neutrophil count (EUR) is independently accounted for by monocytes, adipocytes, and B cell regulatory elements. Then, we annotated SNPs genome-wide using a probabilistic OR gate as follows: 1 - the product of IMPACT scores across these three annotations for SNP $j$. We created 38 composite cell type annotations and observed that these annotations captured significantly more overall enrichment (one-sided paired t-test $P < 2.9e-9$), significantly more per-SNP heritability in terms of $\tau^*$ (one-sided paired t-test $P < 2.0e-7$), and significantly more heritability in the top 5% of SNPs (one-sided paired t-test $P < 1.7e-3$) (**SF17**).

#### Concordance in regulatory basis of complex traits

Not only did we observe shared regulatory biology between populations, but also among traits. Despite weak genetic correlation among different traits, we observed strong correlations of IMPACT annotation $\tau^*$ among traits, revealing large regulatory modules of immunity, white blood cell regulation, red blood cell (RBC) regulation, and body height (**SF18**). These results suggest that while causal effects and variants may differ among traits, there may still be a strongly shared regulatory basis.

<u>Trends of multi-ethnic marginal effect size correlation at various *P* value thresholds</u>

Considering each of the 21 traits, such significant differences were observed at an average of 12 of 17 GWAS *P* value thresholds. Overall, we observed that at lenient *P* value thresholds, the difference in correlation between EUR and EAS effect sizes is more pronounced using IMPACT annotations, suggesting that they may be more effective for prioritizing causal variation particularly when statistical evidence is weak (**SF19**). For example, at the very lenient *P* value threshold of $P < 0.32$, we observed more dramatic improvements in correlation using IMPACT: 5.0x in asthma, 1.1x in RA, 2.8x in MCV, 39.0x in PrCa, and 2.4x in height; all difference of means *P* values $< 0.01$. On the other hand, at more stringent *P* value thresholds, IMPACT annotations offer less of an improvement in multi-ethnic effect size correlation. For example, at a strict *P* value threshold of $P < 3.2e\text{-}7$, we observed modest improvements in correlation using IMPACT: 1.05x in asthma (heterogeneity $P = 0.36$), 1.13x in RA (heterogeneity $P < 7.4e\text{-}3$), 1.12x in MCV (heterogeneity $P < 8.4e\text{-}6$), 1.02x in PrCa (heterogeneity $P = 0.29$), 1.15x in height (heterogeneity $P < 2.6e\text{-}6$).

**Supplement References**

1. Finucane, H. K. *et al.* Partitioning heritability by functional annotation using genome-wide association summary statistics. *Nat. Genet.* **47**, 1228–1235 (2015).
